# Synaptic mechanisms underlying the network state-dependent recruitment of VIP-expressing interneuron-specific interneurons in the CA1 hippocampus

**DOI:** 10.1101/433136

**Authors:** Xiao Luo, Alexandre Guet-McCreight, Vincent Villette, Ruggiero Francavilla, Beatrice Marino, Simon Chamberland, Frances K Skinner, Lisa Topolnik

## Abstract

In the hippocampus, a highly specialized population of vasoactive intestinal peptide (VIP)-expressing interneuron-specific (IS) inhibitory cells provides local circuit disinhibition via preferential innervation of different types of GABAergic interneurons. While disinhibition can be critical in modulating network activity and different forms of hippocampal learning, the synaptic and integrative properties of IS cells and their recruitment during network oscillations remain unknown. Using a combination of patch-clamp recordings, photostimulation, computational modelling as well as recordings of network oscillations simultaneously with two-photon Ca^2+^-imaging in awake mice *in vivo*, we identified synaptic mechanisms that can control the firing of IS cells, and explored their impact on the cell recruitment during theta oscillations and sharp-wave-associated ripples. We found that IS cells fire spikes in response to both the Schaffer collateral and the temporoammonic pathway activation. Moreover, integrating their intrinsic and synaptic properties into computational models predicted recruitment of these cells during the rising to peak phases of theta oscillations and during ripples depending on inhibitory contributions. *In vivo* Ca^2+^-imaging in awake mice confirmed in part the theoretical predictions, revealing a significant speed modulation of IS cells and their preferential albeit delayed recruitment during theta-run epochs, with firing at the rising phase to peak of the theta cycle. However, it also uncovered that IS cells are not activated during ripples. Thus, given the preferential theta-modulated firing of IS cells in awake hippocampus, we postulate that these cells may be important for information gating during spatial navigation and memory encoding.

## INTRODUCTION

Spatial learning, formation of new memories and associated emotional context are encoded by the hippocampus. This structure is populated by a high diversity of local-circuit and long-range projecting GABAergic neurons, which play highly specialized roles in coordinating different computations carried out by principal cells (PCs) and in synchronizing their activity within specific temporal domains [1–3]. This is achieved through the cell type-specific and activity-dependent recruitment of different types of GABAergic neurons during distinct patterns of hippocampal network oscillations [4–7]. The mechanisms that may determine such network behaviour of GABAergic cells have been the focus of intense investigation. From one side, the target-specific properties of the excitatory synapses that different types of inhibitory neurons receive may explain their distinct engagement during specific phases of network oscillations [8–12]. Alternatively, specific inhibitory mechanisms may exist to coordinate selectively the activity of GABAergic neurons [13].

Indeed, in the CA1 hippocampus, the so-called interneuron-specific (IS) inhibitory interneurons are unique in providing synaptic inhibition exclusively to GABAergic neurons. The IS cells that co-express vasoactive intestinal peptide (VIP) and calretinin (CR) and have been classified as type 3 IS cells (IS3s) [14–16] innervate preferentially the somatostatin-expressing (SOM+) *oriens lacunosum-moleculare* (OLM) interneurons [17, 18]. The OLM cells are known to fire at the trough of theta oscillation, but can be silent during the sharp wave-associated ripples (SWRs) via yet unknown mechanisms [4–7, 19]. By inhibiting distal and disinhibiting proximal dendrites of CA1 PCs, OLM cells have been considered essential for modulating the integration of the temporoammonic (TA) *vs* Schaffer collaterals (SC) inputs [20–23], burst firing of PCs [24, 25] and generation of theta oscillations *in vitro* [26–28] and *in vivo* in the ventral hippocampus [29]. Thus, given that synchronous activation of IS3 cells is capable of pacing the OLM interneuron activity at theta frequency [18], the IS3 input to OLM cells may act as a gear mechanism providing for rhythmic gating of the SC *vs* TA inputs in the CA1 area. Whether the IS3 cells are recruited during different patterns of network oscillations *in vivo*, including the theta rhythm, and what can be the underlying synaptic mechanisms operating in these cells, remains however unknown.

As the majority of IS3 cells extends their dendrites through CA1 *stratum radiatum* (RAD) to *stratum lacunosum moleculare* (SLM) [14, 17, 18] and may, therefore, integrate the SC and TA inputs converging within these layers, we examined the SC *vs* TA input integration by IS3 cells as well as the contribution of these inputs to the IS3 cell recruitment during network oscillations in computational models and through recordings in awake mice. We found that IS3 cells can be equally well recruited via either SC or TA inputs due to the synapse-specific transmission properties. Moreover, these cells can preferentially fire during the rising phase/peak of the theta oscillation but are silent during SWRs. Taken together, these data show that, while recruitment of IS cells to theta oscillations during animal locomotion is largely determined by their synaptic properties, additional inhibitory mechanisms may prevent their activation during ripples.

## Results

### Synaptic properties of IS3 interneurons

We took advantage of two transgenic mouse lines (*Vip*-eGFP and *Vip*^Cre^) that express enhanced green fluorescent protein (eGFP) or Cre recombinase in VIP-expressing (VIP+) interneurons (Figure 1A). The *Vip*^Cre^ mice were crossed with a reporter Ai9 line to achieve the tdTomato expression in VIP+ cells [30]. Both mouse lines (*Vip*-eGFP and *Vip*^Cre^) have been previously characterized in details and validated for the expression of the reporter fluorescent proteins selectively in cells expressing VIP endogenously [18, 31, 32]. Consistent with previous findings on VIP expression in hippocampal interneurons [14, 15], in both mouse lines, fluorescently-labeled cells were located throughout the CA1, with many of these having a vertical orientation that resembled the IS3 cell morphology (Figure 1A) [17, 18]. Indeed, 48.8% of VIP+ interneurons in the CA1 area (375/683 cells, n = 3 *Vip*-eGFP mice; 319/739 cells, n = 2 *Vip*-tdTomato mice) were co-expressing CR (Figure S1), consistent with the IS3 cell neurochemical profile [14, 15]. The remainder were VIP+/CR– cells, including a sparsely distributed population of cholecystokinin-co-expressing (CCK+) basket cells [oriens/alveus: 15.2%, pyramidal layer (PYR): 10.2%, RAD: 25.8%, LM: 22.2% of total VIP+ interneuron number/layer; average data for two mouse lines; Figure S1] [33]. We characterized the morphological and electrophysiological properties of VIP+ cells by making targeted patch-clamp recordings from the CA1 RAD VIP+ interneurons in slices from both mouse lines. The biocytin-filled VIP+ cells included in this analysis (n = 72) had somata located predominantly in the RAD, vertical orientation of dendrites extending to the LM and an axon arborizing within the oriens/alveus (O/A) (Figure 1H), consistent with the IS3 cell morphology [14, 15, 17, 18]. These cells had a high input resistance, slow membrane time constant, broad action potentials (APs) and regularly spiking or stuttering firing pattern (Figure S2). The IS3 cell membrane and morphological properties were similar between the two mouse lines (Figure S2), and both lines were used for studying their synaptic properties.

**Figure 1.**
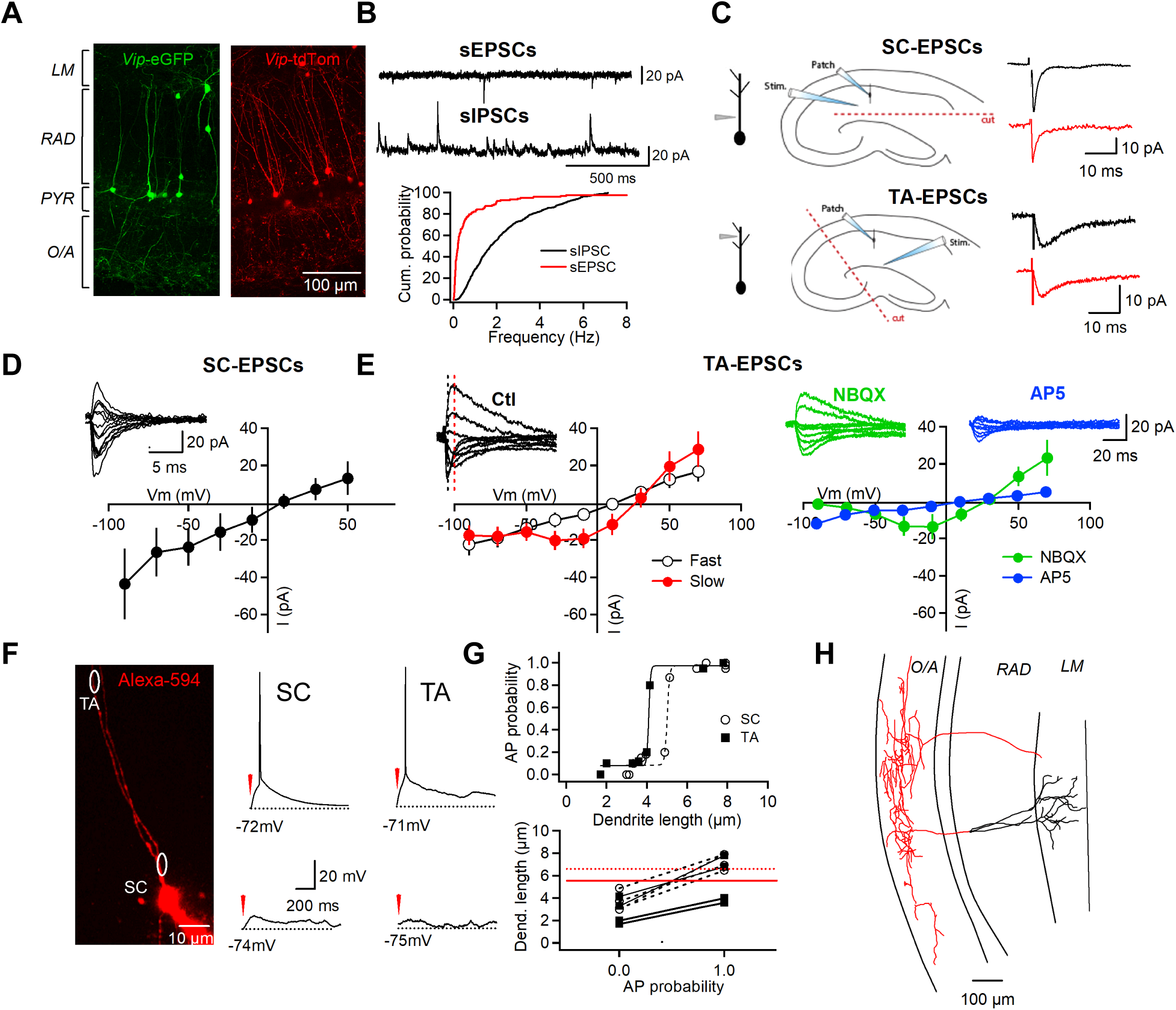
Synapse-specific properties and integration of excitatory inputs in IS3 cells. See also Figures S1, S2, S3 and S4. (A) Two-photon images of CA1 area in slices from *Vip*-eGFP (green) and *Vip*-tdTomato (red) mice. (B) Examples of recordings of sEPSCS and sIPSCs (top) and a summary cumulative distribution for sEPSC (n = 6) and sIPSC (n = 13) frequency (bottom). (C) Left, schematics illustrating the location of the stimulating electrode for SC (top) and TA (bottom) pathway activation; middle, schematics showing the location of surgical cut and stimulating/recording electrodes in slices with surgically isolated SC (top) and TA (bottom) inputs; right, example traces of EPSCs recorded at –70 mV in intact slices (black) and after cut (red). (D) Example traces of SC-EPSCs recorded at different voltage levels and summary I-V curve (n = 5). (E) Left, example traces of TA-EPSCs obtained at different voltage levels, with black and red vertical dotted lines indicating the peak levels for fast and slow EPSC components, respectively, and I-V curves of TA-EPSC fast (black) and slow (red) components (n = 13). Right, example traces of TA-EPSCs recorded in the presence of the AMPAR blocker NBQX (green) or the NMDAR antagonist AP5 (blue) with corresponding summary I-V relationships for the NMDAR (n = 7) and AMPAR (n = 10) components. (F) Left, two-photon image (single focal plane) showing the IS3 cell filled with Alexa-594 during recording and the location of the target areas in proximal (SC) and distal (TA) dendrites for two-photon glutamate uncaging. Right, Representative responses of IS3s when glutamate was released at shorter (bottom, 2-4 *μ*m) and longer (top, 5-8 *μ*m) dendritic areas at SC *vs* TA inputs. (G) Top, representative example of the AP probability in relation to photostimulation extend at two inputs. Bottom, summary data for a group of cells illustrating the transition to AP initiation when the length of stimulated dendrite was increased. Circles correspond to the SC and squares - to the TA inputs, respectively. Red lines indicate the mean threshold for dendritic length, at which it was possible to induce spikes at SC (dotted line) *vs* TA (continuous line) inputs. The difference in the dendritic length was not significant between the two inputs (*P* = 0.173; one-way ANOVA.) (H) Example of an IS3 cell filled with biocytin in 300-µm slice and reconstructed in Neurolucida.

Spontaneous excitatory synaptic drive was very low in these cells (Figure 1B), with an average frequency of spontaneous excitatory postsynaptic currents (EPSCs) of 0.07 ± 0.01 Hz and the average amplitude of 19.9 ± 2.5 pA (n = 6). In contrast, the frequency of spontaneous inhibitory postsynaptic currents (IPSCs) was significantly higher (0.77 ± 0.03 Hz, *P* = 0.002; one-way ANOVA; average amplitude of IPSCs: 19.6 ± 0.8 pA, n = 13), indicating that under basal conditions in slices the IS3 cells receive a dominant inhibitory drive.

The laminar location and frequent monopolar orientation of IS3 cell dendrites (Figure 1H) suggests that these cells may be preferentially driven via the SC and TA inputs from CA3 pyramidal cells and entorhinal cortex, respectively. Accordingly, electrical stimulation in the RAD to activate the SC input elicited kinetically fast and large amplitude EPSCs in IS3 cells (amplitude: 24.0 ± 3.8 pA, rise time: 1.0 ± 0.3 ms, decay time constant: 3.9 ± 1.0 ms; n = 14, Figure 1C–1D), which exhibited a linear current-voltage (*I-V*) relationship (Figure 1D), in line with a major role of the GluA2-containing Ca^2+^-impermeable AMPARs at SC-IS3 cell synapses. In contrast, stimulation within the LM to activate the TA input evoked slower and smaller amplitude EPSCs in IS3 cells (Figure 1C, 1E; Figure S3; EPSC amplitude: *P* = 0.035, rise time: *P* = 0.003, decay time: *P* = 0.012, n = 9 for TA-EPSCs, n = 14 for SC-EPSCs; one-way ANOVA). Importantly, evoked EPSCs exhibited similar input-specific properties in intact slices and in slices with surgical isolation of SC and TA inputs by micro-cuts at the LM or RAD level, respectively (Figure 1C; Figure S3) [34], pointing to the isolated activation of SC and TA pathways in intact slices with our stimulation protocol. Depolarizing the membrane above –50 mV revealed a second slow component to the TA-EPSC (Figure 1E), indicating that NMDA receptors (NMDARs) may be present at TA-IS3 cell synapses. Indeed, the *I-V* relationship of the fast EPSC component was linear whereas that of the slow component was outwardly rectifying (Figure 1E, left; n = 13), suggesting that NMDARs contribute to the TA pathway transmission. Blocking the AMPA receptors with NBQX (12.5 *μ*M) removed the fast component (peak EPSC amplitude decrease to 13 ± 5% of control at –70 mV, n = 10; Figure 1E, right), while application of AP5 (100 *μ*M), an NMDAR antagonist, inhibited the slow component of TA-EPSCs (peak EPSC amplitude decrease to 23.2 ± 5.9% of control at –30 mV, n = 7; Figure 1E, right). Together, these data point to the synapse-specific contribution of NMDARs in IS3 cells, in line with previous reports on the input-specific organization of excitation in other interneuron types [9, 10, 35, 36].

We next explored the spatial integration of SC and TA inputs by IS3 cells using two-photon glutamate uncaging along the IS3 dendrite to mimic the activation of different numbers of excitatory synapses (Figure 1F). In hippocampal slices from *Vip*-eGFP mice, two-photon uncaging (730 nm, 9 ms) of MNI-glutamate (5 mM) along the IS3 dendrite (2–8 *μ*m length) elicited excitatory postsynaptic potentials (EPSPs) (Figure 1F) that were completely abolished by the combination of AMPA and NMDA receptor antagonists (NBQX, 12.5 *μ*M; AP5, 100 *μ*M; n = 3, not shown). We then explored how many synapses were potentially required for triggering action potentials (APs) at SC *vs* TA inputs by uncaging glutamate on proximal (SC input, < 50 *μ*m from the soma, n = 6) *vs* distal (TA input, > 100 *μ*m from the soma, n = 5) dendritic segments of the increasing length. The data showed no difference in the dendritic length necessary to evoke an AP within proximal and distal dendritic segments (SC-IS3: 6.5 ± 0.1 *μ*m; TA-IS3: 5.1 ± 0.1 *μ*m; Figure 1F, 1G; *P* = 0.173, one-way ANOVA). Using the anatomical data on the excitatory synapse density in CR+ dendrites [37], we calculated that 5–7 synapses will be sufficient for triggering an AP at SC and 4–5 synapses at TA-IS3 inputs due to the layer-specific synaptic densities (RAD: 8.5–10 synapses/10 *μ*m; LM: 7.5 synapses/10 *μ*m). Collectively, these results indicate that, when activated synchronously, clusters of ∼5 synapses at different dendritic locations will drive the IS3 cell firing.

We also found that both inputs showed facilitation in response to repetitive electrical stimulation at 10 to 40 Hz (Figure S4A–S4C). However, the facilitation degree at TA-IS3 synapses was significantly higher than that at SC-IS3 inputs for all frequencies tested (10 Hz: *P* = 0.027; 20 Hz: *P* = 0.003; 40 Hz: *P* = 0.0001; n = 18, one-way ANOVA). Moreover, repetitive stimulation at 20–40 Hz resulted in strong summation of EPSPs and increased cell firing (Figure S4A, S4B, S4D). These results indicate that IS3 cells can be driven via both SC and TA inputs during high frequency activity patterns. What can be the inhibitory inputs converging onto the IS3 cells? Morphological studies indicate that hippocampal VIP+ and CR+ cells make synaptic contact with each other [14–16]. To explore the possibility that IS3 cells receive inhibitory inputs from other VIP+ and/or CR+ cells, we injected the Cre-dependent viral vector to express the hM4Di (Gi) designer receptor exclusively activated by designer drug (DREADD) in the CA1 VIP+ or CR+ cells using *Vip*^Cre^ or *Calb2*^Cre^ mice, respectively. Subsequently, sIPSCs were recorded in IS3 cells in control and after application of Clozapine N-oxide (CNO; 10 *μ*M) to silence selectively VIP+ or CR+ CA1 interneurons. The data showed that silencing either cell type resulted in a small but significant reduction in the frequency of sIPSCs in IS3 cells (*VIP*^Cre^;hM4Di: n = 6, *P* = 0.0003, Figure 2A–B; *Calb2*^Cre^;hM4Di: n = 8, *P* = 0.0007, Kolmogorov–Smirnov test; Figure 2C–D), indicating that IS3 cells receive inhibitory inputs from VIP+ and CR+ interneurons. Indeed, we visualized the VIP+, CR+ and VIP+/CR+ axonal boutons on IS3 cell somata (Figure 2E, top; 2F, top) and proximal dendrites (Figure 2E, bottom; 2F, bottom). To examine whether IS3 cells may contact each other, we performed paired recordings between VIP+ cells in slices from *Vip*-eGFP mice. The results showed that IS3 cells were connected to each other via dendritically located synapses, with unitary IPSCs having a small amplitude (19.7 ± 5.5 pA) and high failure rate (80.3% ± 5.5%, n = 4 connected pairs/33 attempts, Figure 2G–H). Taken together, these data indicate that IS3 cells receive inhibitory inputs from other VIP+ and CR+ interneurons [e.g., CR+ type 1 IS (IS1) and VIP+ type 2 IS (IS2) cells), and are connected with each other.

**Figure 2.**
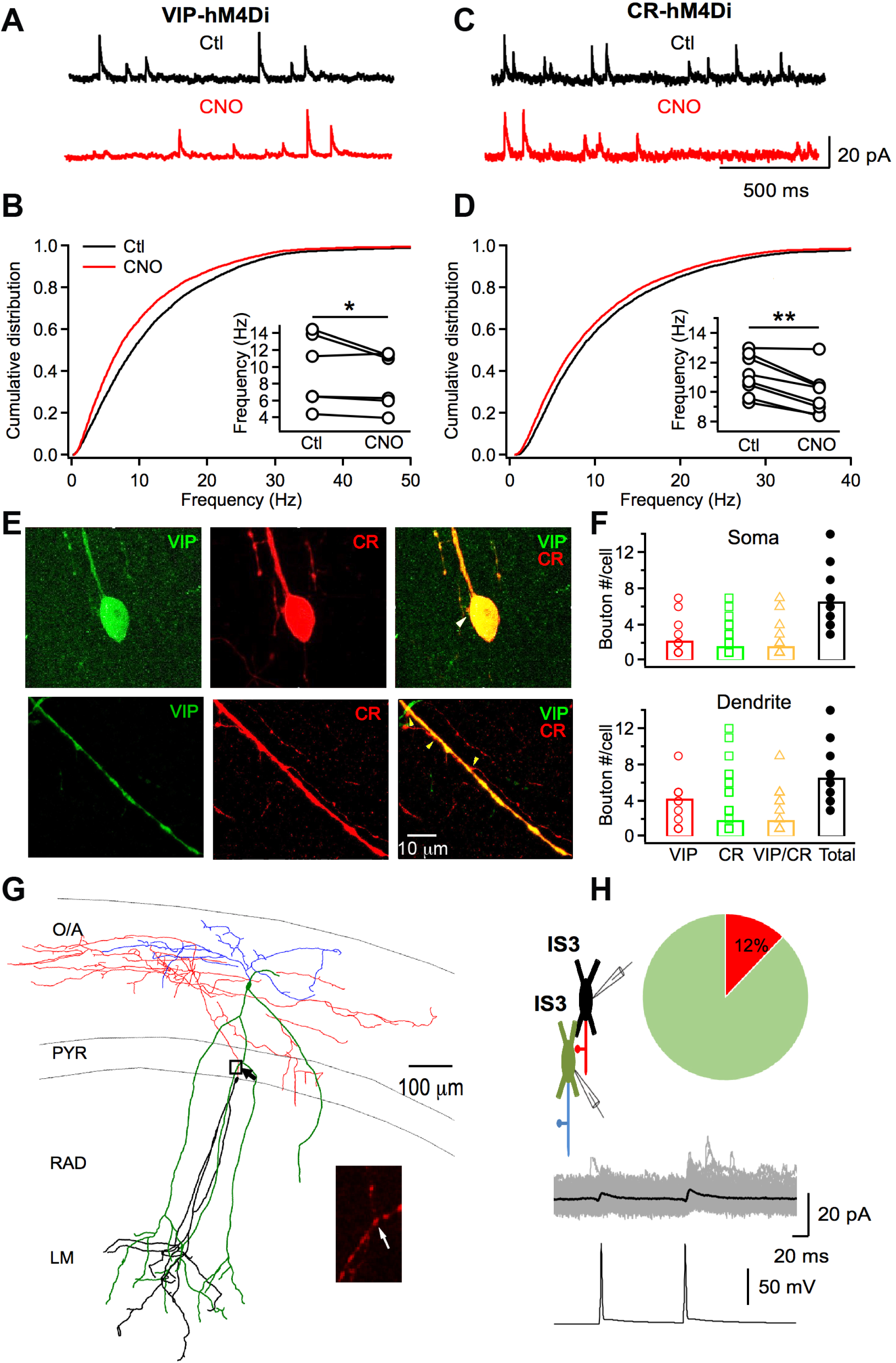
IS3 cells receive inhibitory inputs from VIP+ and CR+ interneurons. (A) Example traces of sIPSCs before (black) and after (red) CNO application recorded in IS3 cells of *Vip*^Cre^ mice expressing a pharmacogenetic silencer hM4Di in the CA1 VIP+ interneurons. (B) Normalized cumulative distribution of sIPSC frequency before (black) and after CNO application (red) in IS3 cells of *Vip*-hM4Di mice. Inset shows mean values obtained in individual cells (n = 6; *P* < 0.05, paired *t* test). (C–D) Same as A and B in *Calb2*^Cre^ mice (n = 8; *P* < 0.01, paired *t* test). (E) Confocal image of the IS3 cell somata (top) and dendrite (bottom) identified based on the co-expression of VIP and CR, that receives VIP+/CR+ (somata) or CR+ (dendrite) axonal boutons indicated with yellow arrowheads. (F) Summary data showing the axonal bouton density (red – VIP+, green – CR+, orange VIP+/CR+ and black – the total number of boutons/cell) contacting the IS3 cell somata (top) and dendrites (bottom). (G) Representative Neurolucida reconstruction (Z-stack from 300-µm slice) of the recorded connected pair of IS3 cells (the axons are shown in red and blue, while dendrites are shown in green and black), with an inset illustrating the axo-dendritic contact from the area indicated with a black square and an arrow in the reconstruction. (H) Schematic of the paired recording between two IS3 cells (top left), a summary pie-chart showing the connection ratio for IS3 cells (top right), and corresponding traces of unitary IPSCs (grey) with the averaged trace (black) in response to presynaptic APs (bottom).

### Synaptic properties of IS3 cells predict their phase-specific firing during network oscillations *in vivo*

To predict the input-specific synaptic recruitment of IS3 cells, we took advantage of the previously developed IS3 cell multi-compartment models [38]. Using experimental data for SC- and TA-EPSCs, as well as IPSCs, we fit the weights, rise and decay times for excitatory and inhibitory synapses onto each dendritic compartment and extrapolated a linear distance-dependent weight rule to generate realistic EPSCs and IPSCs across the entire dendritic arbor of the IS3 cell model (Figure S5A). The weights used here predicted a biologically realistic range of receptors per synapse (Figure S5B), and an increase in the measured reversal potential with distance from soma (Figure S5C), which was in line with experimental observations (Figure 1D, 1E). Using these realistic synaptic parameters, we simulated the *in vivo*-like conditions in IS3 cell model bombarded with synaptic inputs [39]. Two IS3 cell models with (SDprox1; Figure 3, 4) and without (SDprox2; Figure S7) dendritic A-type potassium channels were tested, since both model variants sufficiently captured the electrophysiology of IS3 cells [38]. We applied the theta- and SWR-timed synaptic inputs to predict the network state-dependent firing of IS3 cells *in vivo*. For theta-timed inputs in an *in vivo*-like scenario (Figure 3), the phasic excitatory and inhibitory inputs spiked once per cycle (Figure 3C), with a small amount of noise in their exact timing to enhance the model recruitment during theta-timed inputs (Figure S6A). The model included different proximal and distal dendritic excitatory and inhibitory inputs based on their relative timing during theta oscillations recorded in the CA1 PYR (Figure 3A, 3B) [2, 40]. We found that both the SC- and TA-inputs could drive the phasic recruitment of the IS3 cell (Figure 3D–3F), with the TA-input spiking the IS3 cell during the rising phase, and the SC-input – near the peak of the theta wave (Figure 3G, 3H; Figure S6). Including proximal and distal inhibitory inputs at different phases of theta (i.e. the peak, the falling phase, and the trough), we found a significant reduction in the IS3 cell recruitment when any of the inhibitory inputs were doubled (Figure 3E). Notably though, this effect was seen less with inhibitory inputs occurring near the falling phase and the trough of the theta cycle. Similar results were obtained with SDprox1 (Figure 3) and SDprox 2 (Figure S7) IS3 cell models and, taken together, suggest that IS3 cells would spike between the rising phase and peak of the theta cycle *in vivo*.

**Figure 3.**
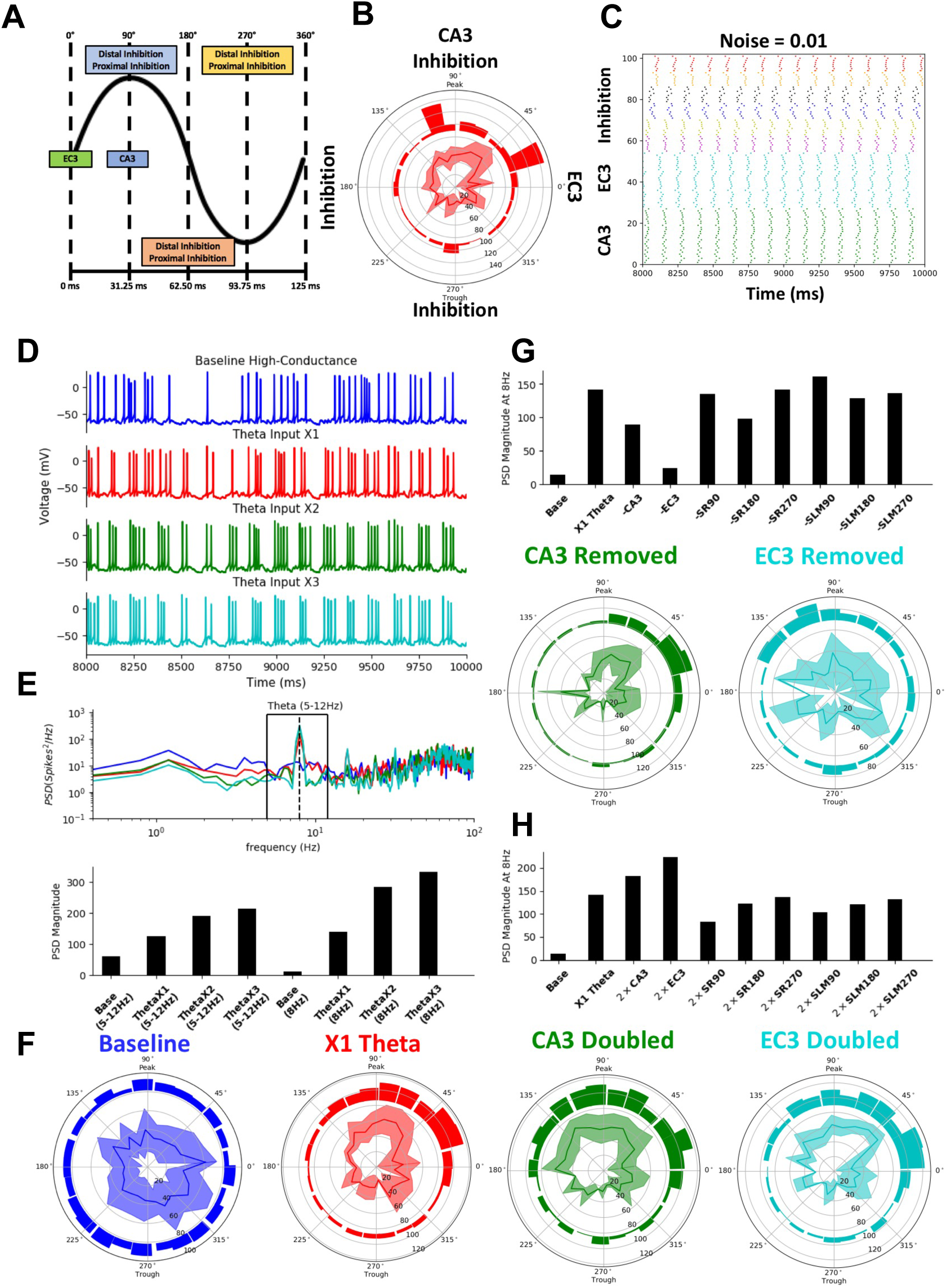
Model predicts firing of IS3 cells during the rising phase/peak of theta wave. See also Figures S5, S6 and S7. (A) Relative timing of different theta-timed presynaptic populations. (B) Same as A, but in polar plot format with an example plot of the model phase preference when theta-inputs are perfectly timed. (C) Example raster plot of the presynaptic theta-timed populations. (D) Example simulated voltage traces of resulting baseline *in vivo*-like conditions, X1 theta inputs, X2 theta inputs and X3 theta inputs. (E) Top subplot: Power spectral density (PSD) of the IS3 cell model spike trains during baseline *in vivo*-like conditions, X1 theta inputs, X2 theta inputs, and X3 theta inputs. Bottom subplot: first four bars show the area under the PSD between 5-12 Hz, and the last four bars show the PSD magnitude at 8 Hz. (F) Polar plots showing phase preference of the IS3 cell model during theta-timed inputs. Inner traces show the mean frequencies of the interspike intervals, binned according to phase (binned standard deviations shown in the shaded areas). Outer histograms show the binned spike phases (bin width = 14.4°). (G) PSD subplots at 8 Hz during removal of specified presynaptic theta-timed populations (top) and theta polar plots (bin width = 14.4°) predicting IS3 phase preference during removal of either CA3 or EC3 (bottom). (H) Same as G, but with doubled presynaptic populations instead of removed.

For the SWR *in vivo*-like scenario [6, 7], we first explored the impact of proximal excitatory inputs mimicking the SC-input from the CA3 PCs during SWR generation (Figure 4A–B, left) [41]. We found that IS3 cells could exhibit an increase in activity above baseline when receiving a SC-input alone (Figure 4C–D). However, including a delayed feed-forward inhibitory input onto the proximal dendrites (RADinh), we found a substantial reduction in the IS3 cell recruitment (Figure 4A–D). Furthermore, including an additional delayed feed-forward inhibitory input onto the distal dendrites (SLMinh, Figure 4A–B, right) did not show any added decrease in the IS3 cell activity (Figure 4C–D). Similar results were obtained with SDprox1 (Figure 4) and SDprox2 (Figure S8) models, and when the synapse location and presynaptic spike timing were re-randomized (Figure 4, 6; Random Seeds 1 *vs* 2). Taken together, these results suggest that IS3 cells can spike during SWRs in response to the CA3 input. However, the activation of feed-forward proximal inhibitory inputs converging onto these cells (e.g. from other VIP+ or CR+ interneuron types) would have a strong dampening effect on their recruitment.

**Figure 4.**
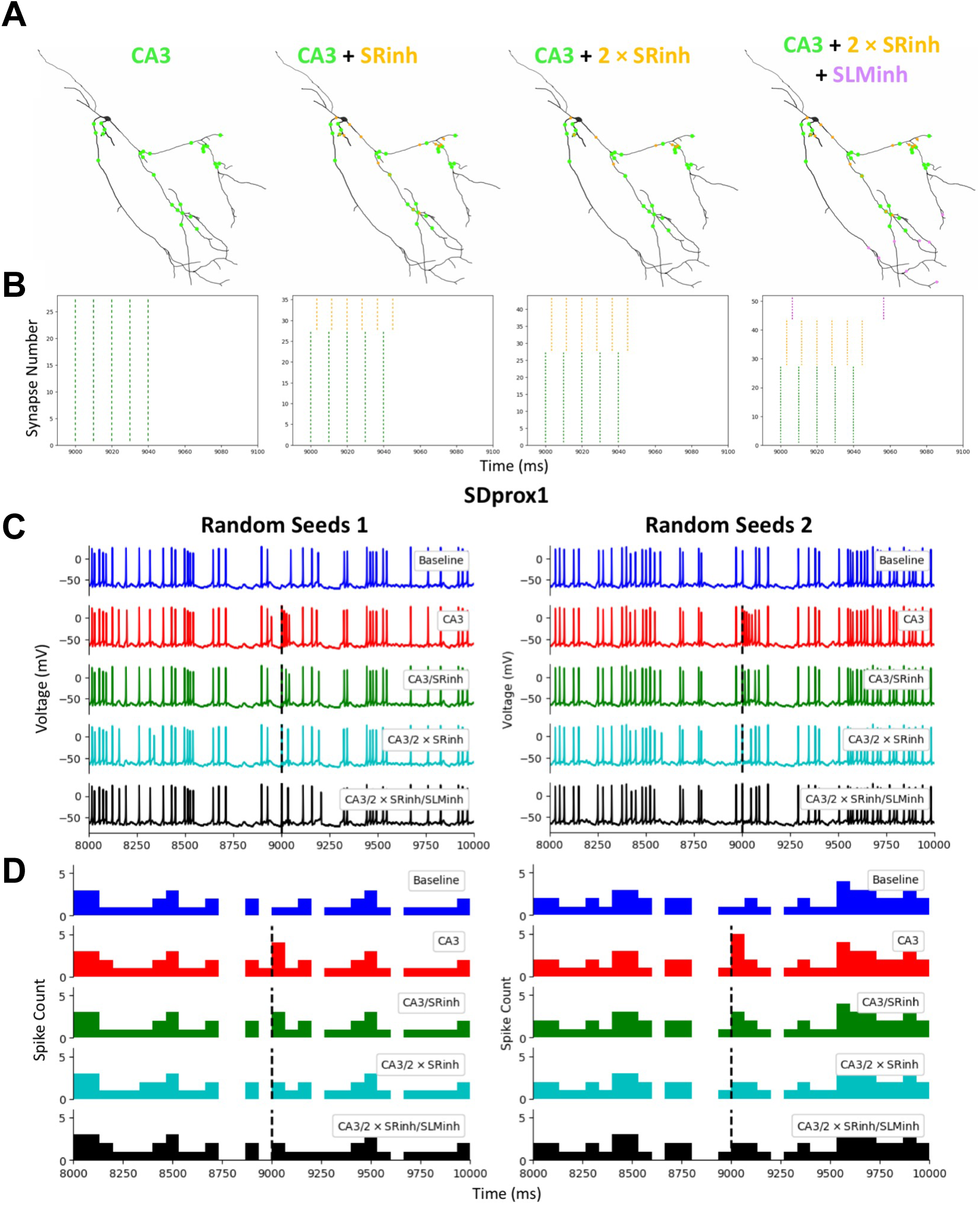
Local inhibitory inputs can dampen the IS3 cell recruitment during SWRs. See also Figure S8. (A) Example synaptic locations of SWR-timed inputs. Note that synaptic locations are chosen randomly in all simulations. (B) Raster plots of SWR-timed presynaptic populations. (C) Example voltage traces during SWR-timed inputs. Dashed black line coincides with the start of the SWR event. Note that we show two model examples where we resample synaptic locations and baseline presynaptic spike times. (D) Peri-stimulus histograms corresponding to the voltage traces shown in C (bin size is 67ms/30 bins). Note that delayed proximal inhibition is sufficient to bring the model spiking back to baseline levels. SRinh - Proximal Feedforward Inhibition, SLMinh - Distal Feedforward Inhibition.

### Delayed recruitment of IS3 cells during theta-run epochs in awake mice

To test the model predictions, we next examined the recruitment of IS3 cells during different network states *in vivo* using two-photon calcium (Ca^2+^) imaging in awake, head-restrained mice trained to run on the treadmill [42]. We expressed a genetically-encoded Ca^2+^ indicator GCaMP6f [43] in VIP+ cells using a Cre-dependent viral vector AAV1.Syn.Flex.GCaMP6f.WPRE.SV40 injected into the CA1 area of *Vip*^Cre^ mice (n = 3 mice). Two-photon somatic Ca^2+^-imaging was performed within the hippocampal CA1 PYR and RAD, the main location site of IS3 cell somata (Figures 1A; S1; 5C), in combination with local field potential (LFP) recordings from the contralateral CA1 PYR (Figure 5D, top) considering that theta oscillations and SWRs are coherent between the two hemispheres [32, 44, 45]. In addition, recordings of the animal’s speed were performed to track locomotion (Figure 5A–B) and analyze hippocampal theta oscillations during theta-run episodes (speed > 1 cm/s; Figure 5E, left) and SWRs during quiet states (speed < 1 cm/s; Figures 5D, bottom and 5E, right) [42, 46].

**Figure 5.**
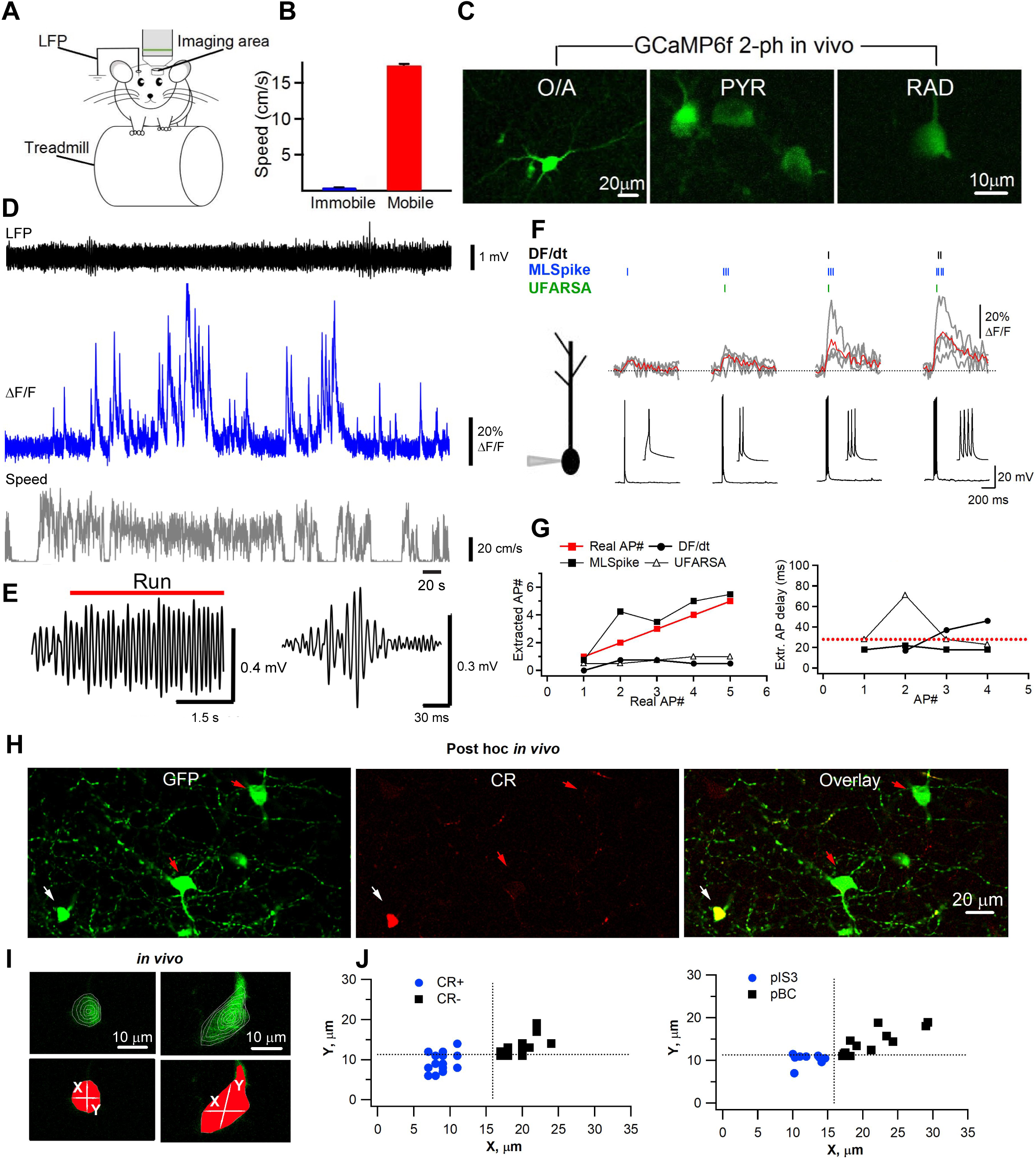
Imaging activity of pIS3 cells in awake mice. See also Figure S9. (A) Schematic of simultaneous Ca^2+^-imaging and LFP recordings in awake mice. (B) Bar graph showing the average running speed during different behavioral states. (C) Representative two-photon images showing VIP+ cells located within O/A, PYR and RAD. Note a smaller size round somata of VIP+ cells within PYR and RAD. (D) Representative traces of simultaneous LFP recording (black) and somatic Ca^2+^ transients (blue) from a pIS3 cell during different behavioral states (grey trace: the higher animal speed during locomotion is reported as a high-frequency step pattern). The raw LFP signal was band-pass filtered in the forward and reverse directions (0.5-1000 Hz, 8^th^order). (E) Expanded theta-run period (left) and ripple event recorded during animal immobility (right). (F) Schematic of patch-clamp recording of GCaMP6f-expressing IS3 cell for *in vitro* calibration experiments (left) and average traces of somatic Ca^2+^ transients evoked in 4 different anatomically confirmed IS3 cells in response to different number of somatically evoked action potentials (right). Red traces correspond to intercell average Ca^2+^ transient. Small vertical lines on top of the Ca^2+^ transient traces indicate the spikes extracted with three different algorithms: DF/dt (black), MLSpike (blue) and UFARSA (green). Note that MLSpike was the most accurate in predicting the spike appearance from somatic Ca^2+^ transients. (G) Left, summary graph showing the relationship between the number of extracted spikes and the number of real spikes. Right, the extracted spike delay as a function of spike number. Both graphs indicate that MLSpike algorithm showed the best performance in detecting the number of spikes and the time of their onset (red dotted line). (H) Representative confocal images of horizontal hippocampal sections processed for *post hoc* immunohistochemical identification following *in vivo* two-photon experiments illustrating the expression of GFP (GCaMP6f, left), CR (middle) and both (right). (I) Two-photon raw (upper) and mask (lower) images of VIP interneurons recorded *in vivo* showing the extraction of somatic X- and Y- parameters for cell identification. (J) Summary data showing the clustering of *in vivo* recorded VIP cells based on the distribution of X- and Y- parameters (pIS3, blue; pBC, black). Dotted lines indicate the cut-off levels obtained from *in vitro* analysis and used for *in vivo* cell segregation (X=15.9 µm, Y=11.3 µm).

To estimate the firing onset in IS3 cells during different phases of network oscillations, we extracted the spike onset times from Ca^2+^ transients (CaTs) using three different algorithms (Figure 5F). These included the Matlab toolboxes MLSpike [47] and UFARSA [48], as well as a custom Matlab code estimating spike times using the speed of the Ca^2+^ signal (DF/Dt) [49]. The algorithms were first fine-tuned for IS3 cell-specific somatic CaTs using *in vitro* patch-clamp recordings of somatic CaTs in response to APs evoked by current pulse injection in anatomically confirmed IS3 cells (Figure 5F, 5G). We found that, among the three algorithms tested, the MLSpike was the most accurate in distinguishing the signal from noise and estimating the spike time onset in IS3 cells (Figure 5F, 5G).

In addition, to identify the IS3 cells within the population of VIP+ cells recorded *in vivo*, including the VIP+ basket cells (BCs), after *in vivo* imaging experiments, the brains were processed for immunolabelling for GFP and CR, the IS3 cell marker (Figure 5H). The results showed that CR– and CR+ cells that were imaged within PYR/RAD separate well into two clusters due to a different soma size (Figure 5I, 5J; CR+: n = 17; CR–: n = 13). In addition, the VIP+/CR– BCs were recognized based on their characteristic axonal pattern within the PYR (Figure 5H, left). These observations were confirmed by analyzing the soma parameters of IS3 cells and BCs derived from the anatomically reconstructed neurons recorded *in vitro* (Figure S9; IS3: n = 36; BC: n = 11). The 3D rendering of somatic surface was used to develop the soma areas and extract the soma diameters in medio-lateral *X*- and rostro-caudal *Y*-dimensions *in vitro* (Figure S9). Based on the data distribution for soma diameter and area in the anatomically confirmed IS3 cells and VIP+ BCs recorded *in vitro* (Figure S9E), the cut-off parameters (*X* ≤ 16 *μ*m, *Y* ≤ 11 *μ*m, soma area size ≤ 152 *μ*m^2^) were established to separate putative IS3 cells (pIS3) from the putative BCs (pBC) *in vivo* (Figure 5I, 5J). We then used these criteria to identify pIS3 and explore their activity during network states associated with locomotion and immobility (Figure 6A–B, n = 18 cells, 221 running periods and 556 stationary states from 2 imaging sessions of 5 min each; see Methods for details). Our data showed that, while the time-varying max theta power was significantly increased during run epochs (Figure 6C), the CaTs were mainly observed during run-stop periods (Figure 6D), pointing to the delayed recruitment of pIS3 cells during theta-run epochs. To further characterize this delay in CaTs, we calculated cross-correlations between CaTs and running speed (Figure 6E) or the time-varying theta power (Figure 6F). Specifically, we looked at cross-correlation magnitudes at zero lags (i.e. zeroth peaks) between CaT, animal speed and LFP as well as their maximal peak magnitudes with corresponding lag times (Figure 6E, 6F; Figure S10A–D).

**Figure 6.**
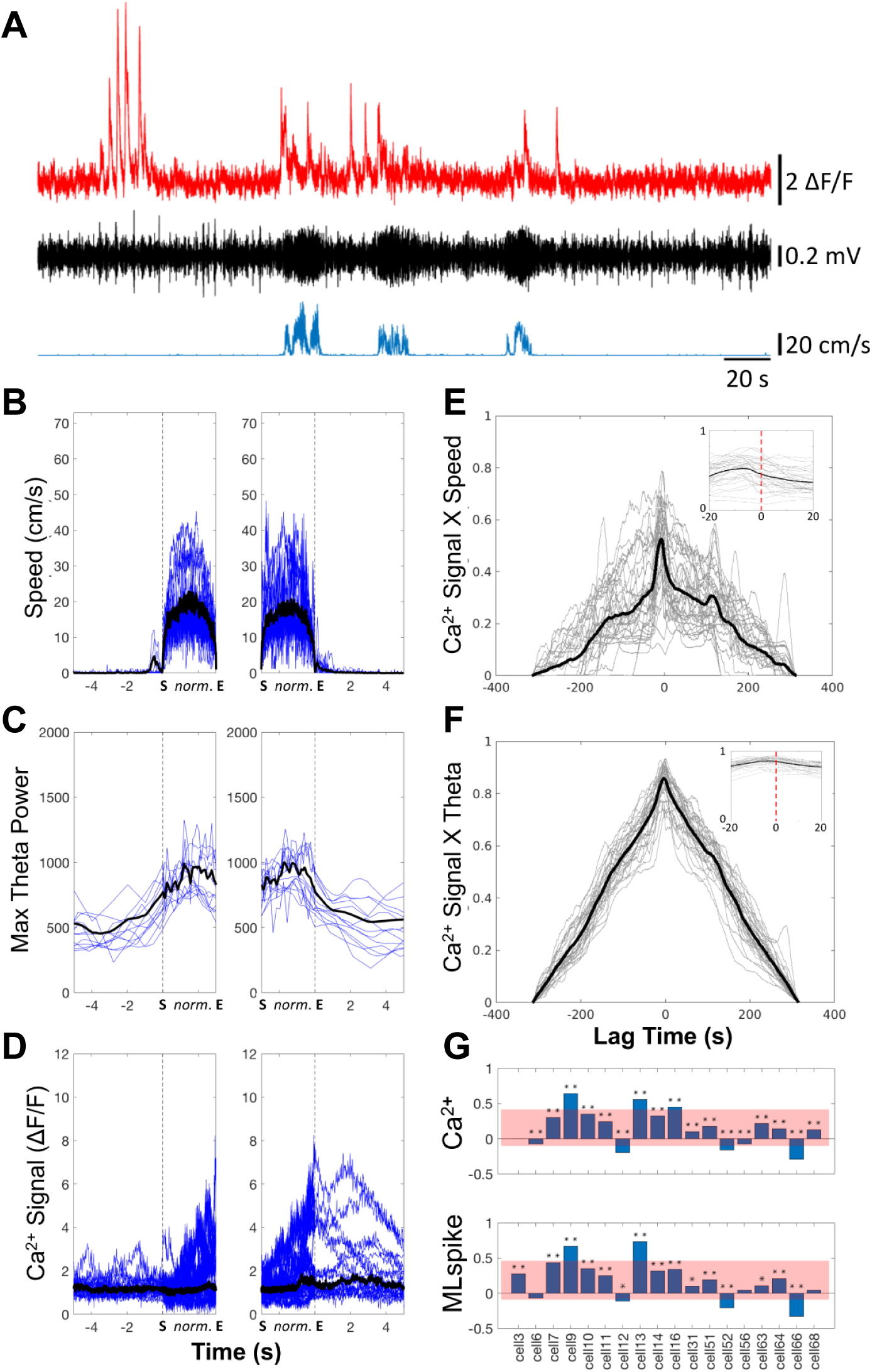
pIS3 cells are active during theta-run epochs. See also Figures S10 and S11. (A) Representative simultaneously recorded traces of a Ca^2+^ signal from a pIS3 cell (red), a theta-filtered (5-12Hz) LFP recording (black), and animal speed (blue trace). (B–D) Average traces of speed (B), theta power (C), and Ca^2+^ signals (D) during run-starts (left panels; averages are across 64 run-starts selected across 17/18 cells) and run-stops (right panels; averages are across 69 run-stops selected across 18/18 cells). The vertical dashed lines show the transition into (left panels) or out of (right panels) run epochs. “S” denotes the start time of a run epoch, and “E” denotes the end time of a run epoch. For the run epoch portions in the time axes, the time vector is normalized (i.e. by using Matlab’s interp1(_) function) such that the run epochs occupy the same span across the x-axis, despite having variable durations. The times outside of these run epochs are shown normally (i.e. ± 5s pre- and post- run epochs). Blue lines show the mean traces for each cell. The mean of means across cells are shown in black, except for the Ca^2+^ signal in D, which shows the median of means. (E–F) Cross-correlations between the Ca^2+^ signal and speed (E), and the time-varying theta power (F). Average traces across all recordings are shown in black, and individual session traces are shown in grey. Speed cross-correlations (E) were quite variable but generally tended to exhibit large peaks near zero. Theta-filtered cross-correlations (F) also exhibited a mean large peak near zero. Inset plots show a zoomed in time scale of -20s to 20s (Y-axes are the same as the larger plots). (G) Pearson correlations of speed with either Ca^2+^ signal magnitude (top subplot), or MLspike estimated spike rates (bottom subplot; \**P* < 0.05; \*\**P* < 0.001). Red area indicates the mean of Pearson correlations ± the standard deviation of Pearson correlations.

Significant zeroth peaks indicated a strong correlation between the CaTs and the running speed (Figure 6E; Figure S10A–B) or CaTs and the theta power (Figure 6F; Figure S10C–D). Similar positive relationships between the CaTs/estimated spikes and the animal speed (Figure 6G) or CaTs/estimated spikes and the theta power (Figure S10E) were obtained using Pearson’s correlations. The lag time of the maximal peaks indicated the approximate delay with which the events from one signal lagged behind the events from another signal. Specifically, negative maximal peak lags indicated that CaTs followed the theta-run epochs. For both, the speed and the theta power, we found that, despite significant correlations between signals, most cells (n = 15/18 for speed; n = 15/18 for theta power) showed negative maximal peak lags (Figure S10A–D), indicating that CaTs were delayed relative to the theta-run epoch onset. Importantly, in line with previous observations [4–7], imaging somatic CaTs in CA1 O/A interneurons, which exhibit consistent phase-locked firing during theta oscillations, showed reliable recruitment of these cells during theta-run epochs, with zero lag between the CaT onset and change in the animal speed (Figure S11), indicating that the delayed recruitment of pIS3 cells was not due to technical artifact. Together, these data indicate that while the activity of pIS3 cells correlates well with running speed and theta oscillation power, these cells exhibit a delayed recruitment.

### IS3 cells prefer to fire near the rising/peak phases of theta oscillations *in vivo*

Given a strong correlation between the CaTs and the theta power (Figure 6F), we next analyzed the theta-phase distribution of IS3 cell spiking, using the MLSpike algorithm for spike extraction and onset time estimates (Figure 7A). In line with significant correlation between the CaTs and the animal speed, the algorithm extracted a larger fraction of spikes during locomotion than during immobility in the majority of cells (Figure 7B), highlighting a higher firing activity of pIS3 cells during locomotion. We then analyzed the theta-phase distribution of spikes during periods of running and high-theta activity. Pooling all of the spike phases from all cells together (Figure 7C), we observed a significant non-uniform distribution (*P* < 1e-11, Rayleigh’s Tests) and a significant polarity towards the circular mean across all spike phases (Table 1; *P* < 1e-12, V-Tests). To characterize the dispersion of this distribution, we analyzed its vector length (vector lengths closer to 1 indicate higher clustering around the mean, i.e. less dispersion) and angular deviation (high values with a maximum of 81.03 indicate a large amount of dispersion). We found a vector length much smaller than 1 (Table 1) and angular deviations close to the maximal possible value of 81.03° (Table 1), indicating a broad distribution of spikes across theta phases. To further investigate this result, we performed the second-order analyses in which the mean of the mean angles across all cells was examined. For spike phase means, we found a significantly non-uniform distribution (*P* < 1e-4, Rayleigh’s Test), as well as a significant mean spike phase preference with most vector lengths towards the rising/peak phase of the theta cycle (*P* < 1e-5, V-Test; Figure 7D). Similar results were obtained when using the UFARSA and DF/dt spike extraction algorithms (Figure S12 and Table 1) and are consistent with the model prediction that modulation from TA and SC pathways cause IS3 cells to fire preferentially during the rising/peak phase of the theta oscillation.

Finally, it has been reported that OLM interneurons, the main postsynaptic target of IS3 cells [18], were often silenced during SWRs [4–7], which, according to our model (Figure 4), could result from the activation of IS3 cells via the SC input. However, the model also predicted that inhibitory inputs converging onto IS3 cells could dampen their recruitment during SWRs (Figure 4C, 4D). Thus, to further expand upon theoretical predictions, we analyzed somatic CaTs recorded in pIS3 cells in relation to SWRs during immobility. Surprisingly, we did not observe any apparent CaTs that coincided with ripple episodes (Figure 7E), indicating that IS3 cells are likely silent during ripples. Indeed, applying the MLSpike algorithm, out of the total number of spikes we could only extract a few in relation to SWRs, and they were all preceding or following these events (Figure 7F, bottom). Moreover, no change in the spike probability was observed during SWRs (Figure 7F, top). Thus, as the model suggests, while IS3 cells can be driven via the SC pathway, activation of feedforward inhibitory inputs converging onto these cells may prevent their recruitment during SWRs.

**Table1.** Circular Statistical Tests.

| Pooled Spike Phases | DF/Dt | UFARSA | MLspike |
| --- | --- | --- | --- |
| Mean Angle | 65.82 | 56.15 | 70.85 |
| Vector Length | 0.06 | 0.12 | 0.12 |
| Angular Deviation | 78.53 | 76.08 | 76.18 |
| Rayleigh's Test (P-Value) | 4.16e-20 | 1.17e-4 | 2.19e-12 |
| Rayleigh's Test (Z-Stat) | 44.58 | 9.03 | 26.76 |
| V-Test (P-Value) | 0.00 | 1.07e-5 | 1.28e-13 |
| V-Test (V-Stat) | 733.80 | 70.34 | 465.54 |
| Mean of Means | DF/Dt | UFARSA | MLspike |
| Mean of Mean Angles | 56.89 | 57.08 | 68.39 |
| Mean of Vector Lengths | 0.12 | 0.31 | 0.17 |
| Angular Deviation of Means | 52.37 | 64.32 | 43.28 |
| Rayleigh's Test (P-Value) | 1.48e-3 | 0.08 | 3.01e-5 |
| Rayleigh's Test (Z-Stat) | 6.10 | 2.46 | 9.19 |
| V-Test (P-Value) | 2.38e-4 | 0.01 | 9.01e-6 |
| V-Test (V-Stat) | 10.48 | 6.66 | 12.86 |

**Table 2.**
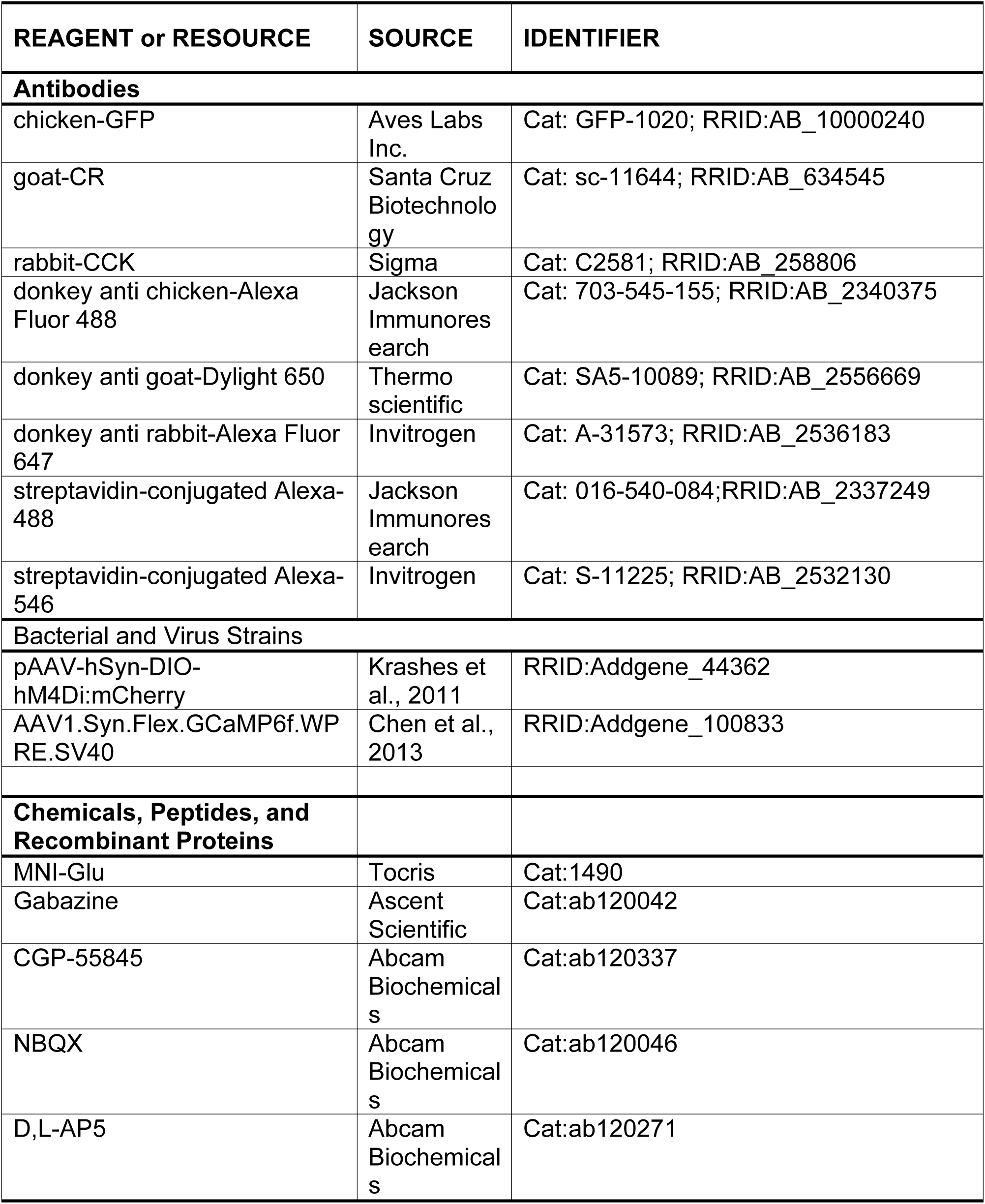
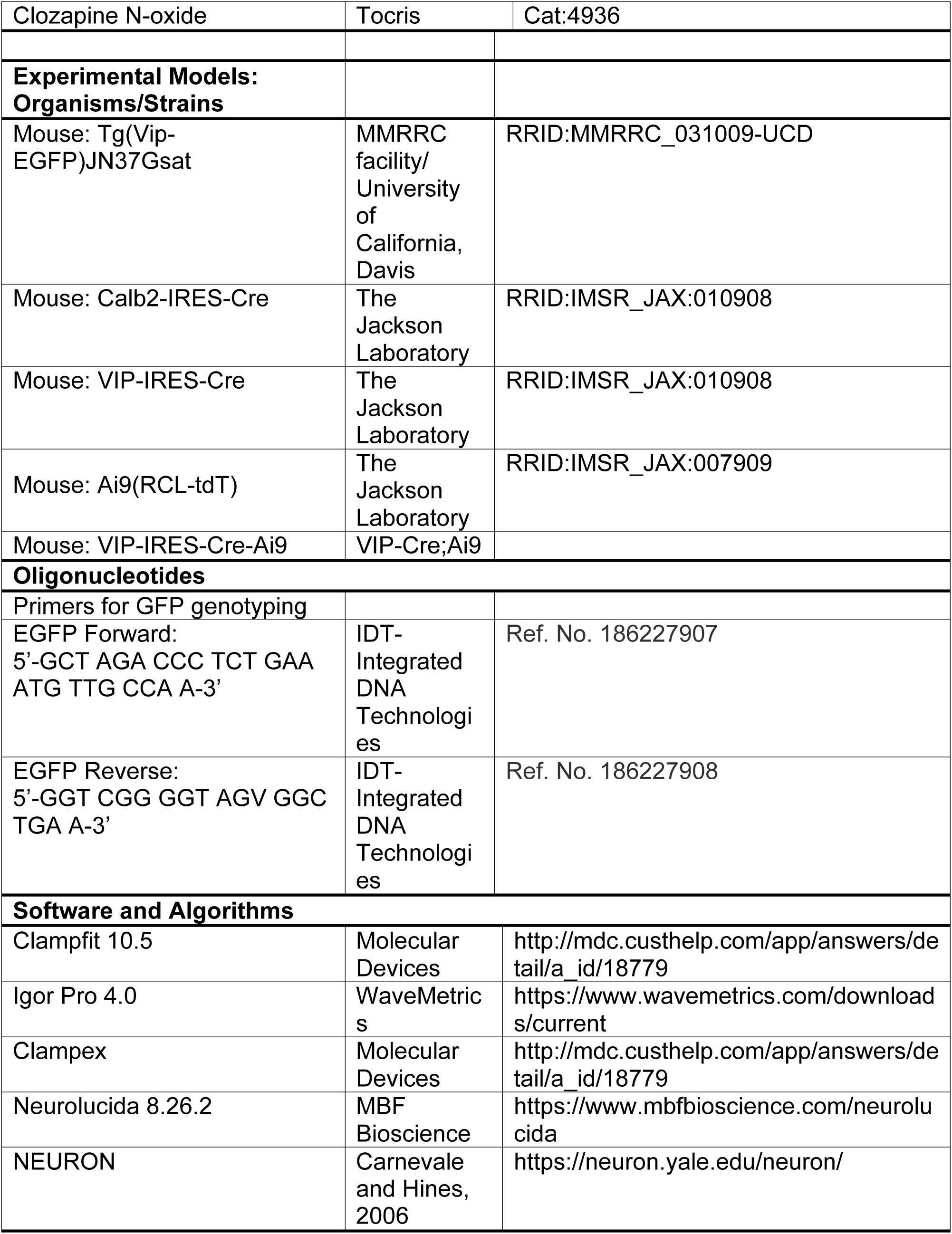
KEY RESOURCES TABLE.

**Figure 7.**
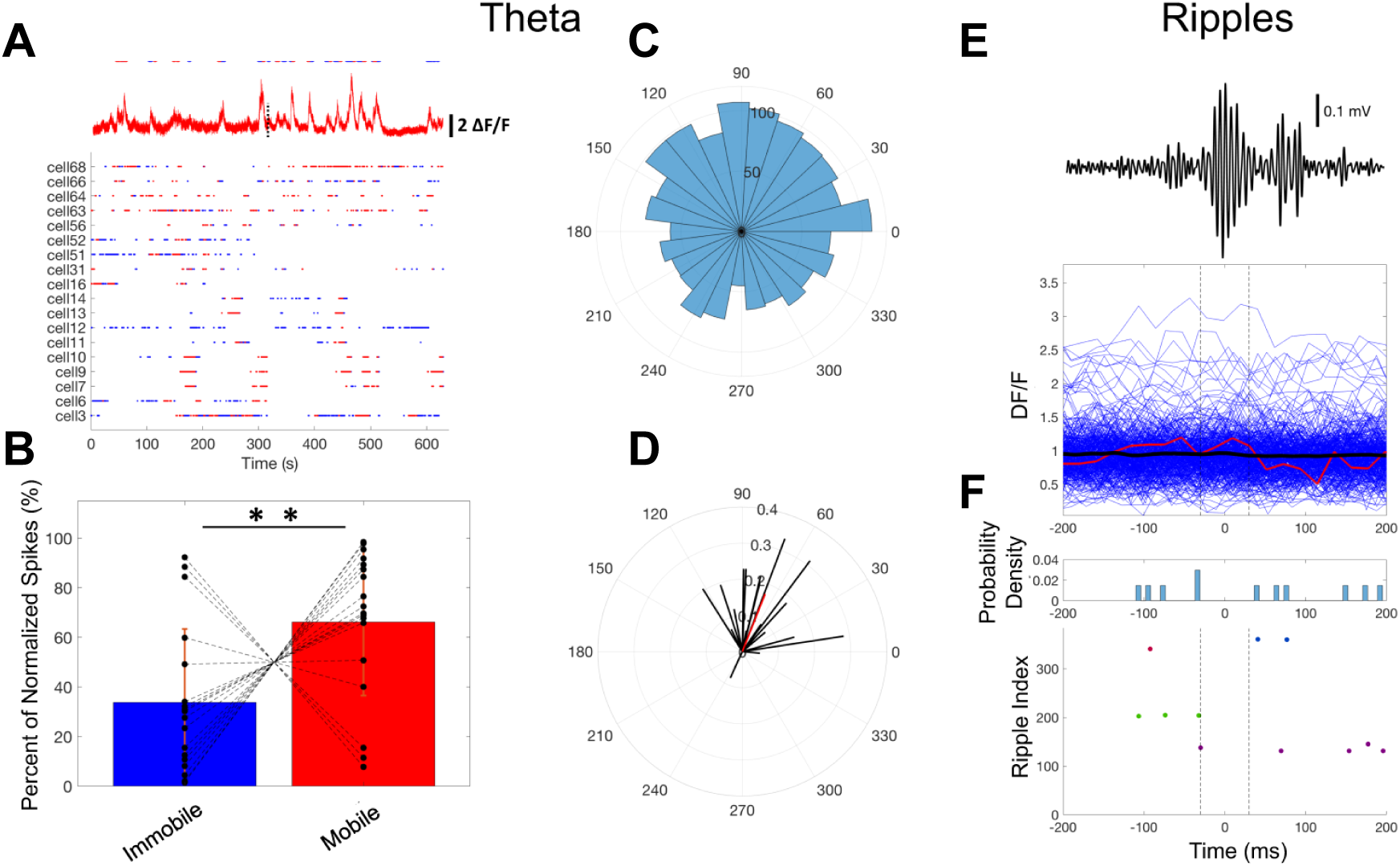
IS3 cells prefer to fire near the rising/peak phases of theta oscillation and are silent during ripples. See also Figure S12. (A) Raster plots for each cell, where blue dots are estimated spikes during immobile states, and red dots are estimated spikes during mobile states. Above the raster plot is a Ca^2+^ trace from cell #66, above which are the estimated spike extractions using the MLspike algorithm. The dashed line in the Ca^2+^ traces indicates the time point where there was a 1 min break between recording sessions. (B) Percent of spikes for each state normalized relative to the states where the animal was spending most of their time (statistical significance across states was evaluated using paired-sample t-tests). The dashed lines indicate which data points belong to the same cell (black dots). Bars indicate the mean, and the error bars indicate the standard deviation (\*\**P* < 0.01). (C) Pooled distribution of spike phases (i.e. across all cells) relative to the theta-filtered LFP, where spike count is shown on the radial axis (bin width = 14.4°). This pooled distribution was significantly non-uniform (Rayleigh’s test), with a significant non-uniform polarity (v-test) towards the circular mean of the distribution (see Table 1). (D) Mean of means test. Each line shows the mean spike phase preference of a cell along the polar axis, where the vector length is shown along the radial axis. The red line indicates the mean of mean angles (i.e. polar axis) and the mean vector length (i.e. radial axis). This distribution was significantly non-uniform (Rayleigh’s test), and had a significant polarity (V-Test) towards the circular mean of the means (see Table 1). (E) pIS3 cells are not active during SWRs. Peri-stimulus time histogram of all Ca^2+^ traces (blue lines) that occur during SWRs. The red line shows the Ca^2+^ trace that corresponds to the example SWR shown above the plot. The black line shows the mean across all Ca^2+^ traces. The dashed lines highlight the ±30 ms time window surrounding the SWR, which is centered at 0 ms along the time axis. (F) Peri-stimulus time histogram of the spike times estimated from the MLspike algorithm. Note that the ripple index on the y-axis highlights how many ripples were analyzed, and the different dot colors corresponds to estimated spike times from different cells. Bin size in top subplots is 8 ms (i.e. 50 bins).

## Discussion

Here, we demonstrate that IS3 cells receive TA and SC excitatory inputs with input-specific glutamate receptor composition and integrative properties. Despite a different dendritic location, both TA and SC-IS3 synapses were able to drive the IS3 cell firing when at least five closely spaced inputs were activated synchronously. Furthermore, these cells receive inhibitory inputs from the VIP+ and CR+ CA1 interneurons, and are synaptically coupled to each other. Computational models of IS3 cells incorporating realistic synaptic conductances predicted their firing during the rising phase/peak of the theta cycle under *in vivo*-like conditions, with the SC and TA inputs providing the largest degree of modulation in terms of the spike timing of IS3 cells. During SWRs, the models predicted that CA3 input alone would be sufficient to recruit these cells but activation of feedforward inhibitory inputs may dampen their activity. Two-photon Ca^2+^ imaging of pIS3 cells in awake mice revealed delayed recruitment of pIS3 cells during theta-run episodes, with a preferential firing at the rising phase/peak of the theta cycle as well as their silence during SWRs. Taken together, these data indicate that, while synaptic properties of IS3 cells determine in large their recruitment to network activity, additional inhibitory mechanisms may control the activity of these cells during specific network states.

The TA *vs* SC pathway-specific synaptic transmission has been well characterized in PCs [50, 51]. Compared to the SC synapses, those formed by the TA input exhibit slower currents and a higher NMDA/AMPA receptor ratio [50]. Similar to PCs, the distribution and subunit composition of postsynaptic glutamate receptors shows the input- and cell type-specific properties in hippocampal interneurons, with a profound impact on the input integration and induction of synaptic plasticity [52–56]. Specifically, the excitatory synaptic transmission at TA and SC pathways was compared in neurogliaform cells [57], with NMDA and AMPA receptors being reported at both synapses. Our data indicate that TA-IS3 EPSCs have slower kinetics than SC-IS3 EPSCs, likely due to dendritic filtering, and are mediated in part by the activation of NMDARs. The AMPAR EPSC components of both TA- and SC-EPSCs showed linear *I-V* relationships, indicating the expression of the Glu2A-containing AMPARs at both pathways. These data are consistent with the transcriptomic analysis of VIP+ neocortical [58] and hippocampal [59] GABAergic neurons, which revealed a highly enriched *Gria2* (GluA2 subunit) mRNAs in VIP+ cells. In addition, the *Grin2b* (GluN2B subunit) and *Grin2d* (GluN2D) transcripts were preferentially expressed in cortical VIP+ neurons [58, 59]. The high NMDA/AMPA receptor ratio controls dendritic spike initiation [60–62], and may facilitate the recruitment of IS3 cells via TA input. Indeed, our data showed that a similar number of synchronously activated SC- or TA-synapses were required for eliciting APs in IS3 cells. It has been documented that the density of excitatory synapses on thick proximal dendrites of CR+ cells is 85.36/100 *μ*m (0.85/*μ*m), while on distal dendrites it is 75.31/100 *μ*m (0.75/*μ*m) [37]. Accordingly, based on the dendritic length required to evoke a single AP in different dendritic microdomains (proximal: ~6.5 *μ*m *vs* distal: ~5.0 *μ*m), the minimal number of synapses leading to the AP generation in IS3 cells may correspond to ~5 on proximal and ~4 synapses on distal dendrites. This observation indicates that additional mechanisms, such as increased NMDA/AMPA receptor ratio [50], synaptic scaling of AMPARs [63] or dendrite site-specific distribution of potassium channels [38] may facilitate distal dendritic integration and spike initiation in IS3 cells.

Regarding the inhibitory inputs, we report that IS3 cells are targeted by the other CA1 IS interneurons, including CR+ (IS1) and VIP+ (IS2) cells, and are connected to each other. These data are consistent with the previous reports on the innervation of VIP+/CR+ cells by VIP+ or CR+ terminals in the rat CA1 hippocampus [15, 16]. However, silencing either CR+ or VIP+ cells resulted in a small albeit significant decrease in the spontaneous inhibitory drive received by the IS3 cells, consistent with a low firing of IS cells under basal *in vitro* conditions due to their hyperpolarized membrane potential [17, 18]. These data indicate that other CA1 interneuron types, such as Schaffer collateral and perforant pathway associated cells, may contact IS3 cells and provide the most of the spontaneous inhibitory drive that these cells receive. In addition, it was shown that long-projecting medial septum GABAergic neurons make multiple synaptic contacts with hippocampal VIP+ cells [64] and may play a role in modulating IS3 cell activity during network oscillations [65–68]. Revealing the cellular identity of all IS3 modulators was beyond the scope of this study and will require new lines of investigation and combinatorial genetic strategies [69].

How can these synaptic mechanisms control the recruitment of IS3 cells during network oscillations *in vivo*? Using computational simulations, we demonstrate the impact of different excitatory and inhibitory inputs on IS3 cell firing. First, the relative timing and level of activity along the SC and TA pathways could define the spike timing of IS3 cells during theta oscillations. Second, removing inhibition of IS3 cells at the falling phase and trough of the theta wave could increase their firing, pointing to the importance of these phase-specific inputs for IS3 cell recruitment. In contrast, doubling inhibition, regardless of the theta phase, would decrease IS3 cell firing, indicating that strong inhibition can hinder the recruitment of these cells. Furthermore, *in vivo* Ca^2+^ imaging of pIS3 cells with estimated spike times confirmed the model predictions on the preferential firing of these cells during the rising phase/peak of the theta cycle, though with a large degree of phasic dispersion. Such high variability in the experimental data relative to the model could result from a number of factors including additional excitatory inputs that were not considered under the current study (e.g. CA1 PCs or subcortical neuronal populations), or cellular diversity within the pIS3 cell population. Indeed, we recently identified a new type of VIP+ long-range projecting (VIP-LRP) cell in the CA1 hippocampus that, in addition to the CA1, innervates subiculum with a region-specific target preference [32]. Further diversity was revealed within the VIP-LRP population, with cells co-expressing the muscarinic receptor 2, CR or proenkephalin in addition to VIP, and having somata located within PYR, RAD or LM. Given that here the soma size of VIP+ cells recorded *in vivo* was used for pIS3 identification and that all CR+/VIP+ cells have a smaller soma size (Figure 5), the subiculum-projecting fraction of this population could be sampled as pIS3 cells and contribute to variability in cellular activity patterns. Indeed, we reported that the subiculum-projecting VIP+ cells are the theta-off cells, as they strongly decrease their activity during theta-run episodes [32]. The latter was seen in a small number of pIS3 cells recorded in the current study (3/18 cells were decreasing activity during theta-run episodes), and may be due to specific modulatory inputs received by the two VIP+ cell types.

The model also predicted that IS3 cells would spike during SWRs driven by CA3 SC input, unless they receive a strong feed-forward inhibition. In line with the latter, we found no correlation between the pIS3 cell activity and SWR appearance *in vivo*. The mechanisms responsible for IS3 cell suppression during SWRs remain currently unclear but, in addition to local and long-range inhibition, may involve subcortical neuromodulatory projections. For example, serotonergic input from the median raphe nucleus was shown to modulate hippocampal SWRs [70], which may depend on the cell type-specific expression of specific types of 5-HT receptors. In addition to a well-documented expression of the excitatory 5-HTR3a by VIP+ cells [71, 72], recent transcriptomic analysis revealed several 5-HT receptor genes involved in the inhibitory signal transduction (e.g., *5Htr1a* and *5Htr1d*) [59]. Whether these candidate receptors are expressed by IS3 cells and lead to suppression in their activity during ripples remains to be explored.

In conclusion, our data reveal the preferential recruitment of IS3 cells specifically during theta oscillations. Thus, a slow inhibition provided by these cells to OLM interneurons via the α5 GABA_A_ receptor-containing synapses [73, 74] may be responsible for pacing the OLM activity at theta frequency and generation of theta-oscillations (e.g., type II theta associated with the emotional context and generated in ventral hippocampus) [29]. Moreover, tuned by specific learning paradigms in relation to animal mood state and experience, such as reward [75] or goal-oriented behavior [76, 77], VIP+ IS cells may play a role of an important modulator of microcircuit activity for information selection and encoding.

## Acknowledgments

This work was supported by the Canadian Institutes of Health Research (CIHR) and the Natural Sciences and Engineering Research Council of Canada (NSERC). Vincent Villette was supported by the PDF fellowship from Savoy Foundation. Alexandre Guet-McCreight was supported by an NSERC CGS-D2 award. We thank Sarah Côté for technical assistance, Dimitry Topolnik for equipment calibration and maintenance, and Armen Saghatelyan for providing *Calb2*-IRES-Cre mice.

## STAR METHODS

## CONTACT FOR REAGENT AND RESOURCE SHARING

Further information and requests for resources and reagents should be directed to and will be fulfilled by the Lead Contact, Lisa Topolnik.

## EXPERIMENTAL MODEL AND SUBJECT DETAILS

Five mouse lines of either sex were used in this study: VIP/enhanced green fluorescent protein (*Vip*-eGFP; RRID:MMRRC_031009-UCD), *Calb2*-IRES-Cre (Jackson #010774; RRID:IMSR_JAX:010774), *Vip*-IRES-Cre (Jackson #010908; RRID:IMSR_JAX:010908), B6.Cg-*Gt(ROSA)26Sor^tm9(CAG-tdTomato)Hze^*/J (Ai9-RCL-tdT) and *Vip*-IRES-Cre-Ai9 (or *Vip*-tdTomato). *Vip*-eGFP line [MMRRC strain #31009, STOCK Tg(Vip-EGFP) 37Gsat], in which eGFP was targeted selectively to VIP+ interneurons (Tyan et al., 2014), was purchased from the MMRRC facility at the University of California (Davis, CA). *Vip*-tdTomato line was generated by breeding *Vip*-IRES-Cre mice with the reporter line B6.Cg-Gt(ROSA)26Sortm9(CAG-tdTomato)Hze/J (Ai9; RRID:IMSR_JAX:007909), stock #007909, The Jackson Laboratory). All experiments were performed in accordance with animal welfare guidelines of the Animal Protection Committee of Université Laval and the Canadian Council on Animal Care. Animals were housed in groups of 2–4 per cage with standard light/dark cycle and *ad libitum* access to food and water. For *in vivo* two-photon Ca^2+^ imaging, the *Vip*-IRES-Cre mice of either sex (P30–60; 25–30 g) were housed separately with standard light/dark cycle and *ad libitum* access to food and water. All animals were randomly assigned to experimental groups.

## METHOD DETAILS

### Patch-clamp recordings in hippocampal slices

Mice (P14-30) were deeply anaesthetized with isoflurane (2%) or ketamine-xylazine (10 mg/ml-1mg/ml). To improve the quality of acute brain slices, mice older than P25 were intracardially perfused with ice-cold sucrose-based artificial cerebrospinal fluid (ACSF) solution containing the following (in mM): 250 sucrose, 2 KCl, 1.25 NaH_2_PO_4_, 26 NaHCO_3_, 7 MgSO_4_, 0.5 CaCl_2_, and 10 glucose (290-310 mOsm/L), aerated constantly with carbogen (95% O_2_/5% CO_2_). Transversal hippocampal slices (300 µm) were obtained using a vibratome (VT1000S; Leica Microsystems or Microm; Fisher Scientific) in ice-cold sucrose-based ACSF and transferred to the recovery solution containing (in mM): 124 NaCl, 2.5 KCl, 1.25 NaH_2_PO_4_, 26 NaHCO_3_, 3 MgSO_4_, 1 CaCl_2_, and 10 glucose (295–300 mOsm/L, pH 7.4) at 33–37 °C, following which they were kept in the same carbogen-aerated solution at room temperature for at least 1 h before recordings. After 1 hour of recovery, slices were transferred to the recording chamber continuously perfused with heated ACSF solution (at 30–32 °C) containing (in mM): 124 NaCl, 2.5 KCl, 1.25 NaH_2_PO_4_, 26 NaHCO_3_, 2 MgSO_4_, 2 CaCl_2_, and 10 glucose (290-310 mOsm/L, pH 7.4). eGFP- and tdTomato-expressing cells were identified using blue and green light illumination, respectively, at upright microscope (Nikon Eclipse FN1) equipped with 40x/0.8N.A objective, and visualized with differential interference contrast for patch-clamp recordings. Two-photon images of eGFP and tdTomato cells in acute slices were acquired using a two-photon microscope (TCS SP5; Leica Microsystems) equipped with a 25x water-immersion objective (NA, 0.95) and coupled to the Ti:Sapphire laser (Chameleon Ultra II; Coherent; > 3W, 140 fs pulses, 80 Hz repetition rate) tuned to 900 for eGFP or to 950 nm for mCherry detection. Borosilicate glass capillaries (1B100F-4; World precision Instruments Inc.) were used to make patch pipettes for whole-cell patch-clamp recordings (tip resistance, 3–6 MΩ). For voltage-clamp recordings, the pipette was filled with a Cs^+^-based solution (in mM): 130 CsMeSO_4_, 2 CsCl, 10 diNa-phosphocreatine, 10 HEPES, 4 ATP-Tris, 0.4 GTP-Tris, 0.3% biocytin, 2 QX-314, 0.1 spermine, pH 7.2–7.3, 280–290 mOsm/L. For current-clamp recordings, a K^+^-based intracellular solution was used, containing (in mM): 130 KMeSO_4_, 2 MgCl_2_, 10 diNa-10 HEPES, 4 ATP-Tris, 0.4 GTP-Tris and 0.3% biocytin (Sigma), pH 7.2–7.3, 280–290 mOsm/L. Passive membrane properties were recorded immediately after cell membrane rupture, active membrane properties were recorded in current-clamp mode in response to somatic current steps (from –200 to +800 pA). In some experiments, the following pharmacological agents were used: gabazine (10 µM, Ascent Scientific, Cat:ab120042), CGP-55845 (2 µM, Abcam Biochemicals, Cat:ab120337), NBQX (12.5 µM, Abcam Biochemicals, Cat:ab120046), D,L-AP5 (100 µM, Abcam Biochemicals, Cat:ab120271) and Clozapine N-oxide (CNO; 10 µM, Tocris, Cat:4936). All drugs were perfused into the bath for at least 10 min before assessing their effect. Spontaneous EPSCs were acquired in voltage clamp at –70 mV in the presence of gabazine. Spontaneous IPSCs were recorded at the reversal potential for EPSCs (0 mV). For electrical stimulation, a bipolar stimulating electrode (tip diameter, 2–3 µm) made from quartz theta glass (O.D.: 1.2 mm; I.D.: 0.9 mm; Sutter instrument) was placed in the CA1 LM or RAD. The EPSCs were evoked by electrical stimulation (1–5 pulses, 5–40 Hz, pulse width 0.2 ms) every 30 seconds in intact slices or in slices with microcut surgical isolation of SC or TA inputs as described in Jarsky et al. (2005). For paired recordings, two *Vip*-eGFP cells were recorded simultaneously: one in current-clamp (presynaptic cell) and the second – in voltage-clamp (postsynaptic cell) mode. The presynaptic IS3 cell was held at –60 mV, while the postsynaptic IS3 cells – at 0 mV to record unitary IPSCs. To study the possible synaptic connection between the two cells, two APs were evoked in the presynaptic cell with brief current injection (0.8–1 nA, 2–3 ms). In case of synaptic connection, short-latency (< 5 ms) unitary IPSCs were detected in the postsynaptic cell. For pharmacogenetic experiments, acute hippocampal slices were prepared from *Vip*-IRES-Cre or *Calb2*-IRES-Cre (CR-Cre) mice expressing hM4Di:mCherry in VIP+ or CR+ cells, respectively. Whole-cell patch-clamp recordings were obtained from vertically oriented mCherry-expressing cells with somata located within CA1 RAD that resembled the morphology of IS3 cells. sIPSCs were recorded at 0 mV before and after CNO application (10 µM). The series resistance (Rser) was monitored throughout the experiments by applying a –5 mV voltage step at the end of each sweep. Recordings with changes in Rser > 15% were discarded from the analysis. Data acquisition (filtered at 2-3 kHz and digitized at 10kHz; Digidata 1440, Molecular Devices, CA, USA) was performed using the Multiclamp 700B amplifier and the Clampex 10.5 software (Molecular Devices). All recorded cells were filled with biocytin and confirmed *post hoc* as IS3 cells. The morphology of selected cells was validated using 3D reconstruction in Neurolucida.

### Two-photon glutamate uncaging

For two-photon glutamate uncaging, Alexa Fluor-594 (20 µM) was included in the intracellular solution to visualize the cell morphology. The caged compound MNI-Glu (5 mM, Tocris, Cat:1490) was applied into the recording chamber. Local photolysis of caged glutamate was achieved by using a two-photon Ti:Sapphire laser tuned to 730 nm (laser power, 25–30 mW) and coupled to Leica TCS SP5 microscope equipped with a 40x (0.8 NA) water-immersion objective. The proximal (< 50 µm from soma) or distal (> 100 µm from soma) dendritic segments of different length (2–8 µm) were illuminated for 9 ms (the smallest photostimulation duration that can be achieved in *xyt*-mode with our two-photon system). The size of the uncaging region was increased gradually until an action potential was evoked (Chamberland et al., 2010). Uncaging-evoked EPSPs or action potentials were recorded in current-clamp mode. To avoid photodamage, the laser power did not exceed 40 mW, and the photostimulation trials were spaced by 30-s rest intervals. In the end of the experiment, two-photon Z-stacks of cell filled with Alexa Fluor-594 were acquired with laser tuned to 800 nm to visualize the cell morphology.

### Stereotaxic injections

The viral vectors AAV1.Syn.Flex.GCaMP6f.WPRE.SV40 (RRID:Addgene_100833) or pAAV-hSyn-DIO-hM4Di:mCherry (Penn Vector Core; RRID:Addgene_44362) were injected in the CA1 area of *Vip*-IRES-Cre or CR-Cre mice to express GCaMP6f or hM4Di:mCherry selectively in VIP+ or CR+ interneurons. Viruses were produced at the University of North Carolina Vector Core facility. For stereotaxic injection, mice were deeply anaesthetized with intraperitoneal injection of ketamine–xylasine mixture (100 /10 mg kg^-1^) and fixed in a stereotaxic frame (Kopf Instruments). Viral vector was delivered using a microprocessor-controlled nanoliter injector (Word Precision Instruments) according to the following coordinates: CA1 (two sites were injected): site 1: AP –2.0 mm, ML +1.6 mm, DV –1.3 mm; site 2: AP, –2.5, ML, +2.1, DV, –1.3 mm. Viral vector was injected with the speed of 1 mm/min. The total volume was 100 nl. The pipette was kept for 5 min after injection and then withdrawn slowly, and the scalp was sutured. Mice were treated with a postoperative pain killer buprenorphine (0.1 mg/kg^−1^; 48 h) for 3 consecutive days or buprenorphine slow release (0.1 mg/kg^-1^) once. In case of AAV1.Syn.Flex.GCaMP6f.WPRE.SV40 injection, animals were allowed to recover for 7 –10 days before the implantation of hippocampal imaging window.

### *In vivo* two-photon Ca^2+^-imaging in awake mice

Two-photon somatic Ca^2+^-imaging of neuronal activity was performed simultaneously with contralateral local field potential (LFP) recording in head-fixed awake mice running on a treadmill. One week after viral injection, a glass-bottomed cannula was inserted on top of the dorsal hippocampus and secured with Kwik-Sil at the tissue interface and Superbond at the skull level [78]. A single tungsten electrode (33 Ω–CM/F, California Fine Wire) for LFP recordings was implanted in the PYR of the contralateral CA1 region and a reference electrode was implanted above the cerebellum [42]. The head plate was oriented medio-laterally at 7–13° using a four-axis micromanipulator (MX10L, Siskiyou) and fixed with several layers of Superbond and dental cement. Mice were allowed to recover for several days with postoperative pain killer treatment for 3 consecutive days (buprenorphine, 0.1 mg/ kg^-1^; 48 h).

To fix the awake animal under the objective of two-photon microscope, the head plate was clamped to a custom-made *X-Y*-moveable metal frame containing a treadmill [42]. The treadmill was equipped with lateral walls to increase animal contentment and coupled with an optical encoder allowing for synchronous acquisition of running speed and LFP signal. For habituation to head-restricted condition, animals were handled and head-fixed for 5–15 min twice per day for 3–5 consecutive days before beginning the experiments. During experiments, animals showed spontaneous alternations between two behavior states: immobility and walking-running periods. The LFP signal and animal running speed were acquired at a sampling frequency of 10 kHz using the DigiData 1440 (Molecular Devices), AM Systems amplifier and the AxoScope software (v10.5, Molecular Devices). Two-photon imaging was performed using a Leica SP5 TCS two-photon system coupled with a Ti:Sapphire laser (Chameleon Ultra II, Coherent), tuned to 900 nm. A long-range water-immersion 25× objective (0.95 NA, 2.5 mm working distance) was used for fluorophore excitation and light collection to external photomultiplier tubes at 12 bits. Image series were acquired at axial resolutions of 2 µm/pixel and temporal resolutions of 48 images/sec. For each cell recorded, two 5-min imaging sessions were acquired. For each animal, the total length of imaging did not exceed 1 h. After imaging session, animals were returned to home cage. Between animals, the treadmill was cleaned with tap water. IS3 cells imaged *in vivo* were identified *post hoc* using morphological and neurochemical (expression of CR) criteria (Figure 5H–5J, Figure S9).

### Immunohistochemistry and morphological analysis

For immunohistochemical analysis of molecular markers expressed in hippocampal CA1 VIP+ cells, animals were intracardially perfused with sucrose-based ACSF followed by 4% paraformaldehyde (PFA) and 20% picric acid in phosphate-buffered saline (PBS), then the brain was extracted and fixed in 4% PFA/picric PBS overnight at 4 °C. On the next day, fixed brains were embedded in 4% agar. Hippocampal sections (thickness, 40–70 µm) were obtained using a vibratome (VT1000; Leica Microsystems or PELCO EasySlicer) and stored in PB sodium azide (0.5mg/ml) solution. Sections were permeabilized with 0.1–0.3% Triton X-100 in PBS and incubated in blocking solution containing 20% normal serum for 1 h. Then sections were incubated with primary antibodies at 4°C for 24–48 h. After that, sections were incubated with conjugated secondary antibodies for 2–4 h, rinsed, and mounted on microscope slides. Confocal images were obtained using a Leica TCS SP5 imaging system or Zeiss slide scanner (Axio Scan.Z1) equipped with a 488-nm argon, a 543-nm HeNe, or a 633-nm HeNe lasers and a 20x (NA 0.8), a 40x (NA 1.4) or a 63x (NA 1.4) oil-immersion objectives. The primary antibodies used were chicken-GFP (1:1000; Aves Labs Inc., Cat: GFP-1020; RRID:AB_10000240), goat-CR (1:1000; Santa Cruz Biotechnology, Cat: sc-11644; RRID:AB_634545), rabbit-CCK (1:800; Sigma, Cat: C2581; RRID:AB_258806). The secondary antibodies used were: donkey anti-chicken Alexa Fluor-488 (1:1000, Jackson Immunoresearch, Cat: 703-545-155), donkey anti-goat Dylight-650 (1:250; Thermo scientific, Cat: SA5-10089), donkey anti-rabbit Alexa Fluor-647 (1:250; Invitrogen, Cat: A31573).

For axonal bouton analysis in IS3 cells, 70-µm slices obtained from the *Vip*-eGFP mice were processed for eGFP and CR immunoreactivity, and IS3 cells were identified based on the co-expression of VIP (eGFP) and CR. Confocal Z-stacks were acquired using a 63x oil-immersion objective with a 0.2-µm step, and the VIP+, CR+ or VIP+/CR+ boutons contacting the IS3 cell somata or dendrite (distance: 10–100 µm from the soma) were identified and counted in every cell. In total, 20 cells from 5 mice were analyzed.

For *post hoc* morphological analysis, patched cells were filled with biocytin (0.6 mg/ml, Sigma) during patch-clamp recording. After recording, 300-µm slices were fixed in 4% PFA at 4°C overnight. To reveal biocytin, the slices were permeabilized with 0.3% Triton X-100 and incubated at 4 °C with streptavidin-conjugated Alexa-488 (1:1000, Jackson Immunoresearch, Cat: 016-540-084) or Alexa-546 (1:1000, Invitrogen, Cat: S11225) in Trizma buffer. Z-stacks of biocytin-filled cells were acquired with a 1.5-µm step and merged for detailed reconstruction in Neurolucida 8.26.2 (MBF Bioscience).

### Computational model

#### Model details and code

The models used in this study (SDprox1 and SDprox2) were previously custom-built to simulate electrophysiological characteristics of IS3 cells (ModelDB accession #: 223031) [38]. Models were simulated in the NEURON environment and results were plotted using customized Python and Matlab scripts. Note that all code and scripts can be accessed online (synaptic optimization code: https://github.com/FKSkinnerLab/IS3-Cell-Model/tree/master/LayerSpecificInputTests; rhythmic input simulations code: https://github.com/FKSkinnerLab/IS3-Cell-Model/tree/master/RhythmTests) [79].

#### Model synapse details

For the synapse model, we used NEURON’s built-in Exp2Syn function, which models synaptic current as a two-state kinetic scheme.

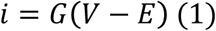

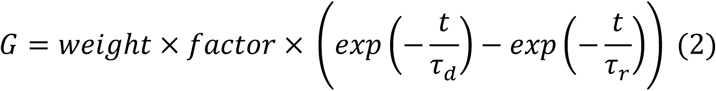

Where *i* is the synaptic current, *G* is the synaptic conductance, *E* is the reversal potential, *V* is the membrane potential, *weight* is the synaptic weight, *factor* is a NEURON process that is used to normalize the peak synaptic conductance to the value of weight, *t* is time, *τ_r_* is the rise time, and *τ_d_* is the decay time.

We used NEURON’s Praxis Optimizer to fit the synaptic parameters for each compartment to an experimental EPSC or IPSC. Excitatory synapses located less than 300 µm from soma were fit to a SC-evoked EPSC, and excitatory synapses located at distances exceeding 300 µm from soma were fit to a TA-evoked EPSC. All inhibitory synapses were fit to a spontaneous IPSC. We set the *weight*, *τ_r_*, and *τ_d_* as the free parameters in the optimization and assumed that all voltage-gated channels were blocked, since sodium and potassium channel blockers were used during the experimental recordings. To remove any driving forces caused by leak conductances, we also set the leak reversal potential to the voltage-clamped holding potential of the model (0 mV when fitting IPSC recordings, and –70 mV when fitting to EPSC recordings). During the optimizations, we used experimental *τ_r_* and *τ_d_* individual trace measurements as seed values (IPSC: *τ_r_* = 0.25 ms and *τ_d_* = 4.14 ms; EPSC*_Proximal_*: *τ_r_*= 0.45 ms and *τ_d_* = 1.41 ms; EPSC*_Distal_*: *τ_r_* = 1.71 ms; *τ_d_* = 5.04 ms) and used parameter bounds based on literature values (AMPA [80–86]; GABAA [86–88]). These chosen parameter ranges and bounds were the following:

### AMPA

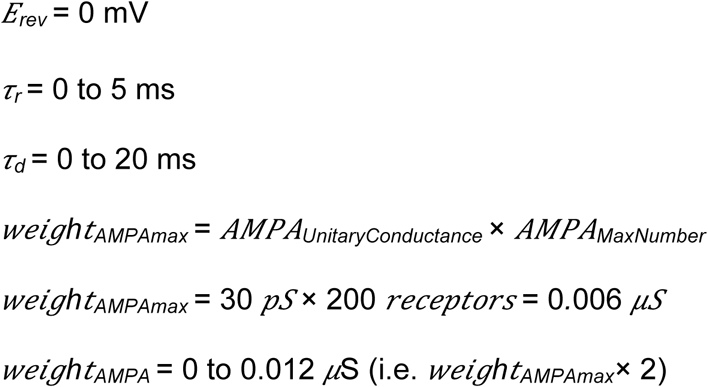

### GABA_A_

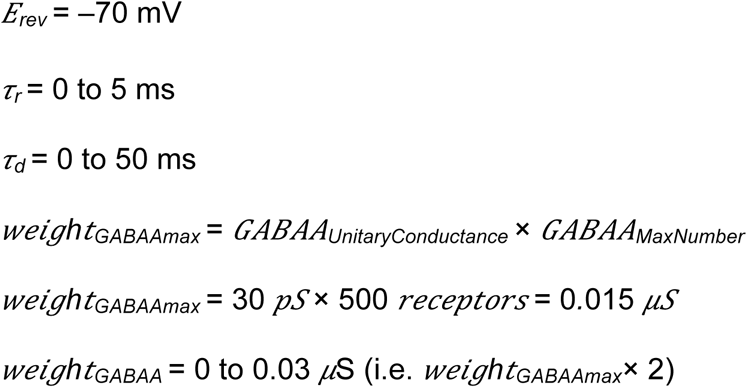

Presynaptic Spike Time = 0 to 6 ms

Using the weight value obtained from the most optimal fit for proximal dendritic synapses as well as the weight value obtained from the most optimal fit for distal dendritic synapses, we applied a linear distance-dependent weight rule for AMPA synapses:

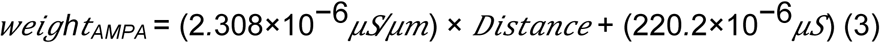

Similarly, using the weight values obtained from the two most optimal fits for inhibitory synapses, we applied a linear distance-dependent weight rule for GABA synapses:

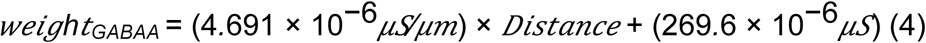

Respectively, the rise times and decay times of all excitatory synapses were fixed to the rise and decay times of their corresponding layer-specific most optimal fits. The rise times and decay times of all inhibitory synapses were fixed to the rise and decay times of the most optimal inhibitory fits: Excitatory RAD: *τ_r_* = 2.9936 × 10^−4^ ms, *τ_d_*= 2.4216 ms; Excitatory LM: *τ_r_* = 6.1871 × 10^−4^ ms, *τ_d_*= 3.1975 ms; Inhibitory: *τ_r_* = 0.1013 ms, *τ_d_*= 4.8216 ms.

To get the ranges of numbers of receptors per synapse for each synaptic location (Figure S5), we computed the following:

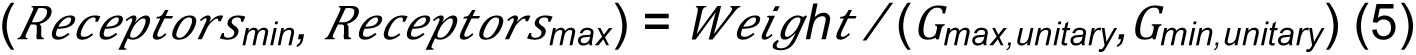

To simulate the experimental protocol for estimating the distance-dependent reversal potential of each excitatory synapse (Figure S5), we held the soma in voltage-clamp mode and incrementally increased the holding potential. The reversal potential of each synapse was recorded as the holding potential when the polarity of the postsynaptic current at the soma reversed.

### Baseline *in vivo*-like Inputs

*In vivo*-like excitatory and inhibitory inputs (i.e. spike rates and number of active synapses) were estimated previously [39]. In all simulations, the baseline *in vivo*-like synaptic locations and presynaptic spike trains were chosen randomly according to a set of random seeds. For SDprox1, the baseline *in vivo*-like inputs have the following parameters: 144 excitatory synapses with 5 Hz rates each, and 8 inhibitory synapses with 50 Hz rates each. For SDprox2, the baseline *in vivo*-like inputs have the following parameters: 144 excitatory synapses with 5 Hz rates each, and 24 inhibitory synapses with 10 Hz rates each.

### Theta-timed and SWR-timed Inputs

For all *in vivo*-like simulations, synaptic locations (i.e. for baseline, theta-timed, and SWR-timed synapses) were chosen randomly, such that synapses were placed anywhere within their designated proximal or distal (or both) dendritic sections [39]. For estimated numbers of X1 theta-timed excitatory inputs, we increased the number of inputs on either proximal or distal dendrites, until a large enough power spectral density (PSD) was obtained (i.e. 50 spikes^2^/Hz). For estimated numbers of X1 theta-timed inhibitory inputs, we first increased a current input at the soma until a base spike rate of 35 Hz was attained. We then increased the number of inhibitory inputs on either proximal or distal dendrites until a large enough PSD was obtained (i.e. 80 spikes^2^/Hz). Between the proximal and distal dendrites for the SDprox1 and SDprox2 models, we estimated approximately 27 synapses per excitatory presynaptic population and approximately 8 synapses per inhibitory presynaptic population. Note that we used these same estimates for SWR-timed presynaptic populations.

Unless otherwise noted, presynaptic spiking for both theta-timed inputs and SWR-timed inputs were idealized, such that they spiked perfectly on time at their prescribed frequencies. For theta, this was 8 Hz for all presynaptic populations. For SWR inputs this varied across different populations (excitatory: 100 Hz, feedforward proximal inhibitory: 120 Hz, feedforward distal inhibitory: 20 Hz; Figure 4B), and the duration was limited to 50 ms.

To simulate both theta-timed and SWR-timed inputs, we used NEURON’s NetStim function to simulate spike trains with corresponding frequencies. For theta-timed inputs, we adjusted the start time of the presynaptic populations depending on their phase preferences. Finally, we also analyzed the effects of fractional randomness on theta-recruitment (Figure S6). The fractional randomness (i.e. noise values between 0 and 1), controls the proportion by which each spike interval is influenced by the set interval value versus a random interval value sampled from a negative exponential distribution with a mean duration of noise × interval.

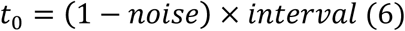

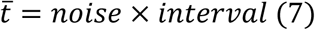

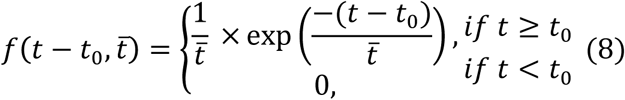

Where *t* is time, *interval* is the set inter-spike interval, and *noise* is the fractional randomness. This probability density function ensures that *t_0_* is the minimum possible interval, that the maximum possible interval is infinite, and that the mean expected interval is (1 − *noise*) × *interval* + *noise* × *interval*. Here we found that theta-recruitment was optimal with a 0.01 level of fractional randomness (noise; Figure S6).

### Data analysis

Data acquired in electrophysiological experiments was analyzed in Clampfit 10.5 (Molecular Devices) and Igor Pro 4.0 (WaveMetrics). For the analysis of the AP properties, the first AP evoked with a current step of +40 to +60 pA within 50 ms from the beginning of the current step was analyzed. The AP amplitude was measured from the threshold to the peak. The AP half-width was measured at the voltage level corresponding to the half of the AP amplitude. Input resistance (R_in_) and capacitance (C_m_) were obtained using a membrane test in Clampex (Molecular Devices). The membrane time constant (*τ*) was measured offline using an exponential fit of voltage response to a hyperpolarizing step of –100 pA. Somata area, dendritic length and dendritic surface were generated automatically from the 3D reconstructions of recorded cells in Neurolucida 8.26.2.

For the analysis of animal behavior on the treadmill during *in vivo* experiments, spontaneous locomotion activity was defined as time points where the speed was greater than 1 cm/s. Immobility periods were defined as the times when the speed was less than 1 cm/s. The run epochs were defined as contiguous periods when the instantaneous speed was larger than 1 cm/s for a minimal time period of 3 s (221 epochs of mean duration 8.95 ± 0.47 s). To identify run-starts for plotting in Figure 6B– 6D, only run epochs where the average speed was below 1 cm/s for the 5 s preceding them were selected (N = 64; 17/18 cells). Likewise, to identify run-stops, only run epochs where the average speed was below 1 cm/s in the 5 s following them were selected (N = 69; 18/18 cells). Stationary epochs were defined as contiguous periods when the instantaneous speed was smaller than 1 cm/s for a minimal time period of 3 s (556 epochs of mean duration 9.31s ± 0.31 s).

The image analysis was performed off-line using Leica LAS, Igor Pro (Wavemetrics, Lake Oswego, USA) and Matlab. For extraction of somatic Ca^2+^-transients, a region of interest was drawn around individual somata to generate the relative fluorescence change (F) versus time trace. For the purposes of automating the analysis, the baseline fluorescence level (F_0_) was taken to be the median of Ca^2+^signal. Somatic Ca^2+^-transients were expressed as ΔF/F = (F - F_0_)/F_0_ or ΔF/F = (F - F_0_)/F_0_ x 100%. This analysis was performed for 18 different pIS3 cells, of which 14 were recorded across two independent imaging sessions (5 min each) separated by 1 min interval, and 4 cells – during one session. Analyses between the two sessions were pooled together for each cell. Cross-correlations were computed using Matlab’s xcorr(—,‘coeff’) function. To extract estimated spike times from Ca^2+^events, we applied three different algorithms: DF/dt, MLSpike and UFARSA. In all cases, we tuned the spike extraction algorithm parameters according to *in vitro* calibration experiments (Figure 5F, 5G). The first method was custom written using Matlab, whereas the other two are previously developed algorithms available in Matlab (see Supplementary Data 1 for details). For the first method, we took the gradient of the Ca^2+^ signal (DF/dt) [49] and chose times where the DF/dt trace had crossed a threshold value of 0.75 times the standard deviation of the DF/dt trace for at least two time-steps. In addition, the Ca^2+^ signal itself needed to be greater its own median plus standard deviation during these same time steps. Secondly, we used the MLspike algorithm [47], which we tuned to be optimal for GCaMP6f data and the levels of noise seen in our recordings. To further optimize the accuracy for each Ca^2+^ trace, we used MLspike’s functions for estimating the noise levels. Compared to the other two methods, MLspike performed best in distinguishing signals from noise and detecting the spikes’ onsets (Figure 5F, 5G). Thirdly, we used the UFARSA algorithm (Ultra-Fast Accurate Reconstruction of Spiking Activity) [48]. While this algorithm provided some spike time estimates, it often under-estimated the number of spikes compared to other algorithms. It is to be noted that large variabilities can exist in the time courses and magnitudes of Ca^2+^transients, even within a given cell type. Therefore, considering all technical challenges associated with spike extraction (i.e. poor time resolution, variability in noise level and Ca^2+^ event shapes), all available methods for spike extraction from Ca^2+^ imaging data cannot be considered as precise, which is why we decided to use three different algorithms to compare the data obtained. Ultimately, these methods were used to provide an estimate of spike phases relative to the theta rhythm, such that they could link back to the model predictions regarding the modulation of IS3 cells *via* CA3 and entorhinal cortex inputs during theta rhythm. To examine somatic Ca^2+^-fluctuations in relation to network oscillations, LFP traces were band-pass filtered in the forward and reverse directions to obtain theta-filtered (5–12 Hz, 4^th^ order) or ripples-filtered (125–250 Hz, 8^th^ order) signals in Matlab as follows: [z, p, g] = butter(order, [min_frequency, max_frequency]/nyquist,’bandpass’); [sos, g] = zp2sos(z, p, g); filtered_signal = filtfilt(sos, g, LFP_signal);

The onset of the theta-run epoch, which was always associated with an increase in theta power, was defined by the beginning of the locomotion period based on the animal speed trace acquired simultaneously with LFP. To examine the time-varying relationships with theta power, we extracted the maximum powers from the spectrograms of the theta-filtered LFP traces between 5 and 12 Hz in Matlab, as follows:

window_theta = 20000;

overlap = [];

freqrange_theta = 0:0.001:20;

[s,f,t] = spectrogram(filtered_signal, window_theta, overlap,freqrange_theta, fs)

a = abs(s);

for l = 1:length(t)

Theta_area(l) = max(a((f>5 & f<12), l));

end

We labeled periods where the time-varying theta power was greater than its mean as periods of “high-theta activity”. For obtaining spike phases from the estimated spike times, we took the Hilbert transform (i.e. Matlab’s hilbert(—) function) of the theta-filtered LFP signal and interpolated the spike phases from this trace. In this case, we set 0° to the rising phase of the theta oscillation.

Ripple events were detected automatically using custom written Matlab code. Specifically, we computed the envelope of the ripple-filtered trace by taking the absolute (abs(—)) of its Hilbert transform (hilbert(—)). We then set a threshold criterion to identify time periods where the envelope was greater than its mean plus 5 times its standard deviation for at least 12.5 ms. We chose this criterion manually to ensure that ripple events were both large enough in amplitude and long enough in duration. These time periods were then marked as ripple periods.

### Quantification and statistical analysis

Data was initially tested for normality using a Kolmogorov–Smirnov or Shapiro-Wilcoxon test. If data was normally distributed, standard parametric statistics such as one-way ANOVA were used (\**P* < 0.05, \*\**P* < 0.01, \*\*\**P* < 0.001) to evaluate statistical significance using SigmaPlot 12.5 (Systat Software, Inc) and Clampfit 10.5 (Molecular Devices). If data was not normally distributed, Dunn’s test or Mann–Whitney test were used for comparisons of multiple groups. The data are presented as means ± SEM. The statistical details can be found in the results, figure legends, and tables. The “n” numbers described in the results indicate the number of cells, unless specified.

Statistical significance of cross-correlations was evaluated by doing a structured reshuffling of 10-s segments of the speed traces or time-varying theta power traces to generate 1,000 surrogate cross-correlation traces. Within this surrogate dataset, the 95^th^ and 99^th^ percentile traces were computed using Matlab’s prctile(—) function, in order to evaluate whether the peak or 0^th^ cross-correlation magnitudes fell above these percentiles. We computed Pearson’s linear correlation coefficient between the different spike rate estimates and Ca^2+^ signals using Matlab’s corr(X, Y, ‘Type’, ‘Pearson’) function. Note that this function was also used to compute Pearson’s linear correlation coefficient between the Ca^2+^ signals and speed traces. For comparisons of percent of normalized numbers of spikes between immobile and mobile states, we used Matlab’s ttest(x, y, ‘Alpha’,) function and paired-sample t-tests. All circular statistical and descriptive measurements were computed using the CircStat toolbox in Matlab. Significance of the non-uniformity of circular spike estimate distributions relative to theta phase were tested using Rayleigh’s method (circ_rtest(—)), and directional circular statistics were calculated using V-Tests (circ_vtest(—)). Mean theta-phase modulation of spike estimates were computed using the circular mean (circ_mean(—)) and measures of dispersion included the vector length (circ_r(—)) and the angular deviation (circ_std(—)).

