## Supplemental Data for "Synaptic mechanisms underlying the network state-dependent recruitment of VIP-expressing interneuron-specific interneurons in the CA1 hippocampus"

**Supplementary Data 1**.
Spike estimation algorithm parameters. Note that only parameters that varied from their default parameter values are included in this table. Parameters and code statements are presented in the same chronological order as how they appear in the code. In particular UFARSA is run twice, the first time being to optimize the noise estimation parameter.

| | |
|---|---|
| **UFARSA** | |
| Initial opt.scale_NoiseSTD | 2.25 |
| opt.remove_drifts | 0 |
| opt.remove_posDeflections | 0 |
| opt.remove_negDeflections | 0 |
| opt.demerging | 1 |
| output1 | [output1,~] = run_UFARSA(opt) |
| Finding peaks | pks = findpeaks(output1.fluors.afterSmoothing) |
| Finding minimum peak | Amp_min = min(pks(pks>(0.5*std(pks)))) |
| Optimized opt.scale_NoiseSTD | Amp_min/output1.std_noise |
| output | [output,~] = run_UFARSA(opt) |
| **MLspike** | |
| pax | pax = spk_autosigma('par') |
| pax.bias | 1 |
| Sigma Estimation | sigmaest = spk_autosigma(catrace,dt,pax) |
| par | par = tps_mlspikes('par') |
| par.a | 0.12 |
| par.tau | 0.76 |
| par.pnonlin | [0.85, -0.0006] |
| par.finetune.sigma | sigmaest |
| par.drift.parameter | 0.08 |
| spikest | [spikest,~,~] = spk_est(catrace,par) |

**Figure Supplements**

**Supplementary Figure 1. Immunolabelling for CR and CCK in the CA1 hippocampal area of *Vip*-eGFP (green) and *Vip*-tdTomato (red) mice.**
(A–B) GFP (green), CR (red) and CCK (red) immunoreactivity in hippocampal CA1 area of *Vip*-eGFP (A) or *Vip*-tdTomato (B) mice. Insets in the lower right corner delineated by a white frame show examples of cells co-expressing GFP and CR or GFP and CCK at a higher magnification.
(C) Summary data for CR and CCK co-expression in the CA1 area of Vip-eGFP and Vip-tdTomato mice.

**Supplementary Figure 2. Membrane and morphological properties of IS3 cells in *Vip*- eGFP (green) and *Vip*-tdTomato (red) mouse lines**.
(A–B) Example traces of voltage responses (top) to depolarizing and hyperpolarizing current steps (bottom).
(C) Comparison of membrane properties and morphological parameters of IS3 cells in *Vip*-eGFP (green) and *Vip*-tdTomato (red) mouse lines. P=0.113 for membrane capacitance.

**Supplementary Figure 3. Comparison of EPSC properties in intact slices and in slices with surgically isolated TA or SC inputs.**
(A–C) Comparison of TA-EPSC (left) and SC-EPSC (right) properties of IS3 cells recorded in intact slices and in slices with surgical microcuts, showing no difference in the EPSC peak amplitude (A), rise time (B) and decay time constant (C).

**Supplementary Figure 4. Input-specific temporal and spatial summation.**
(A) Schematics illustrating the stimulating electrode location for the SC pathway activation (upper), and the example traces in response to repetitive stimulation (10, 20 and 40 Hz) in voltage (middle) and current (bottom) clamp mode.
(B) Same as panel A for TA-stimulation.
(C) Normalized EPSC amplitudes in response to repetitive stimulation of SC *vs* TA inputs. Solid lines and closed symbols indicate TA-EPSC, dashed lines and open symbols indicate SC-EPSC (*10 Hz: $P = 0.0027$, 20 Hz: $P = 0.003$, 40 Hz: $P = 0.0001$; one-way ANOVA. The EPSCs induced by the last three pulses were compared between TA and SC pathway).
(D) The average number of APs generated in response to different stimulation frequencies at TA *vs* SC synapses. The statistical analysis was performed on the APs induced by the fifth pulse at 20Hz. (*$P = 0.01$; one-way ANOVA).

**Supplementary Figure 5. Simulation of synaptic inputs along the model dendritic tree.**
(A) Simulated somatically-recorded EPSCs (blue) and IPSCs (red) generated from synapses at selected locations along the dendritic arbor of the IS3 cell model (i.e. with linearly rectified weights). Note a distance-dependent broadening and amplitude deterioration of both the IPSCs and EPSCs. Red arrows denote the approximate synaptic locations. Dashed brown line shows the border between proximal and distal dendrites at 300 µm from the soma. Labels on the y-axis denote the dendritic subtree labels. RAD – Stratum Radiatum; LM – Stratum Lacunosum Moleculare; AIS – axon initial segment.
(B) Predicted ranges of receptors per synapse, calculated based on the fitted weights (blue) and the linearly rectified distance-dependent weights (red). Note that each *X*-axis data point shows a range, which is larger or smaller depending on the thickness (along the *Y*-axis) of the plotted line. Dashed red line denotes the maximal limits as per previous findings from other cell types in the literature. Line bifurcations in this plot correspond to bifurcations in the dendritic morphology.
(C) Left, Estimated reversal potential of each synapse along the dendritic arbor of the model using a simulated protocol that is similar to what is used experimentally. Note that the true reversal potential of each excitatory synapse is 0 mV, and does not change

during these simulations. Line bifurcations in this plot correspond to bifurcations in the dendritic morphology. Middle–Right, Electrotonic distance (i.e. decay of a 1 mV signal) for voltage flowing into the soma (left) and voltage flowing out of the soma (right). Note that the sub-tree organization approximately follows the same organization as in C, suggesting that reversal potential recordings in IS3 cells are sensitive to distance-dependent voltage decay.

**Supplementary Figure 6. Effect of fractional randomness (noise) on theta-recruitment in the SDprox1 model**.
(A) Example X1 theta raster plots of the presynaptic theta-timed populations at different levels of synaptic noise.
(B) Power spectral density (PSD) and the area under the PSD for 5–12 Hz (first four bars) and at 8 Hz theta at noise levels indicated in A.
(C–E) Polar plots showing the phase preference of the IS3 cell model for X1 (C), X2 (D), and X3 (E) theta inputs (bin width = 14.4°).

**Supplementary Figure 7. SDProx2 model predictions for IS3 cell firing during theta oscillations.** The same as Figure 3D–F using the SDProx2 IS3 model.

**Supplementary Figure 8. SDProx2 model predictions for IS3 cell firing during SWRs.** The same as Figure 4C–D using the SDProx2 IS3 model.

**Supplementary Figure 9. Morphological criteria used for the VIP cell identification.**
(A–B) Neurolucida 3D reconstruction (A) and confocal images (B; maximal projections of 10 sections) illustrating the analysis of somatic parameters in anatomically confirmed IS3 cells filled during *in vitro* patch-clamp recordings with biocytin. 3D rendering of somatic surface was used to derive somatic diameters in medio-lateral (X-axis) and rostro-caudal (Y-axis) dimensions (B).
(C–D) The same as illustrated in A–B but for BCs.
(E) Summary bar graphs illustrating the distributions of somatic diameters in a group of cells (IS3, blue, n = 36; BC, black, n = 11).

**Supplementary Figure 10. Summary statistics of cross-correlations and Pearson correlations between theta power and calcium signal.**
(A–B) Analysis of speed x calcium signal cross-correlations. Dots indicate the lag time of the maximum peaks versus the magnitude of the maximum (A) or zeroth (B) peaks seen in the cross-correlations. Histograms along the ordinate axes show the distributions of cross-correlation maximum peak amplitudes and zeroth peak amplitudes. The color of dots indicates the significance of the crosscorrelation using a 10s structured reshuffling surrogate analysis. Black dots indicate p-value > 0.05, blue dots indicate p-value < 0.05, and red dots indicate p-value < 0.01.
(C–D) Same as A–B but for analysis of theta power x calcium signal cross-correlations.
(E) Same as in Figure 6G but with time-varying theta power instead of speed.

**Supplementary Figure 11. Somatic $Ca^{2+}$ transients in CA1 O/A interneurons of awake mice.**

(A) Two-photon image (single focal plan) of CA1 O/A interneurons expressing GCaMP6f.
(B) Examples of calcium traces (black) obtained from cells indicated with numbers in A (cells 1–5) during different patterns of animal behavior. Red trace shows the animal speed with a dotted line illustrating the locomotion threshold. Red arrowheads point to calcium events occurring during animal locomotion periods.
(C) Correlation plot illustrating the cross-correlations between calcium transients and animal speed for 5 cells illustrated in A (black traces) and an average cross-correlation function for all cells (red). Note a tight correlation between the calcium transient onset and change in the animal speed.

**Supplementary Figure 12. Analysis of *in vivo* estimated spike times data using the DF/Dt and UFARSA methods.**
(A) Percent of spikes detected using DF/Dt (left) and UFARSA (right) algorithms for each state normalized relative to the states where the animal was spending most of their time (statistical significance across states was evaluated using paired-sample t-tests). The dashed lines indicate which data points belong to the same cell (black dots). Bars indicate the mean, and the error bars indicate the standard deviation (**$p < 0.01$).
(B) Pooled distribution of spike phases (i.e. across all cells) relative to the theta-filtered LFP detected with two algorithms, where spike count is shown on the radial axis (bin width = 14.4°). This pooled distribution was significantly non-uniform (Rayleigh’s test), with a significant non-uniform polarity (v-test) towards the circular mean of the distribution (see Table 1).
(C) Mean of means test for spikes extracted with two algorithms. Each line shows the mean spike phase preference of a cell along the polar axis, where the vector length is shown along the radial axis. The red line indicates the mean of mean angles (i.e. polar axis) and the mean vector length (i.e. radial axis). This distribution was significantly non-uniform (Rayleigh’s test), and had a significant polarity (V-Test) towards the circular mean of the means (see Table 1).
(D) Peri-stimulus time histogram of the spike times estimated from the DF/Dt (left) and UFARSA (right) algorithms. Note that the ripple index on the y-axis highlights how many ripples were analyzed, and the different dot colors corresponds to estimated spike times from different cells. Bin size in top subplots is 8 ms (i.e. 50 bins).

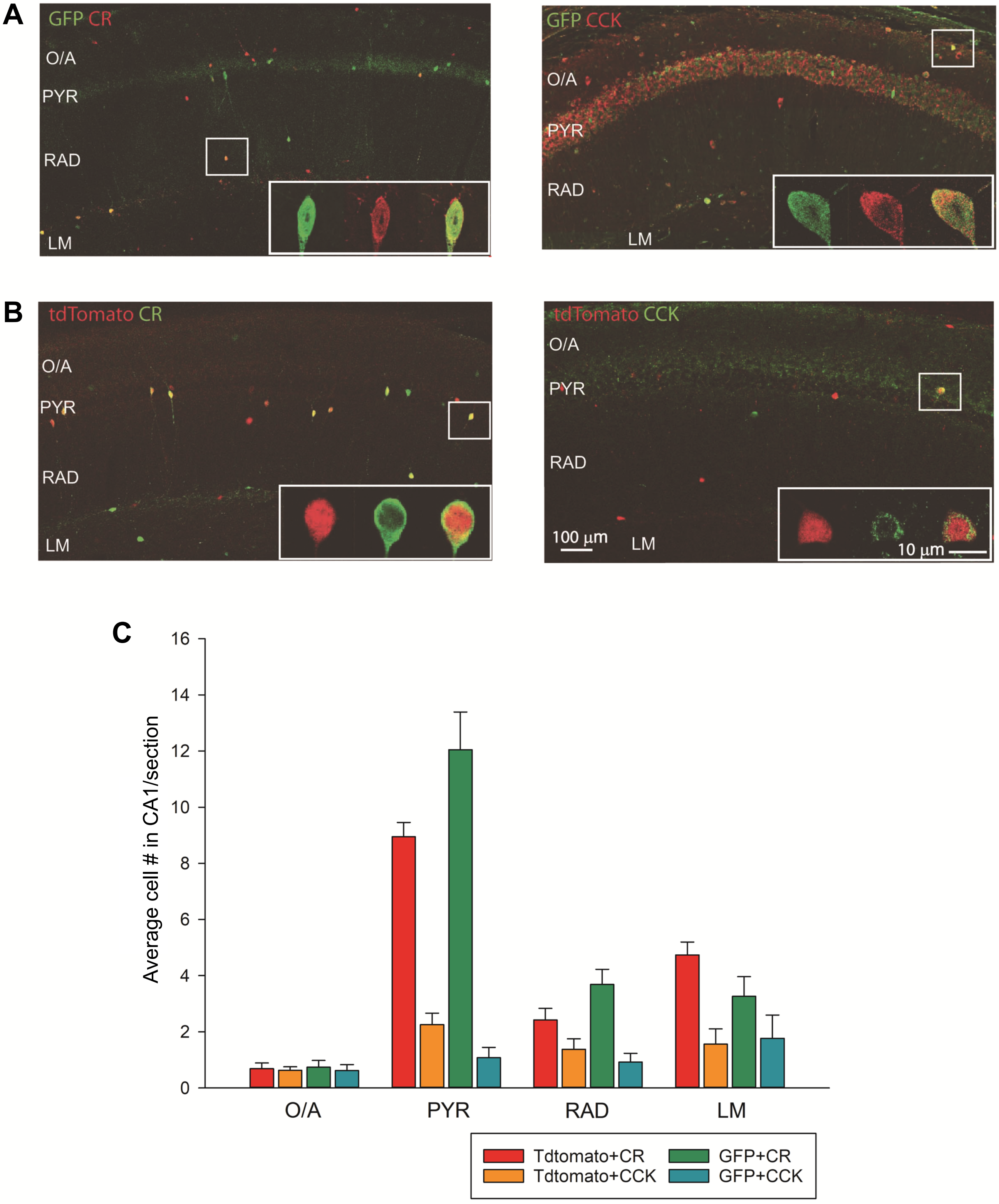

A
GFP CR
O/A
PYR
RAD
LM
GFP CCK
O/A
PYR
RAD
LM
B
tdTomato CR
O/A
PYR
RAD
LM
tdTomato CCK
O/A
PYR
RAD
100 µm
LM
10 µm
C
Average cell # in CA1/section
16
14
12
10
8
6
4
2
0
O/A
PYR
RAD
LM
Tdtomato+CR
Tdtomato+CCK
GFP+CR
GFP+CCK


**Figure S1.**

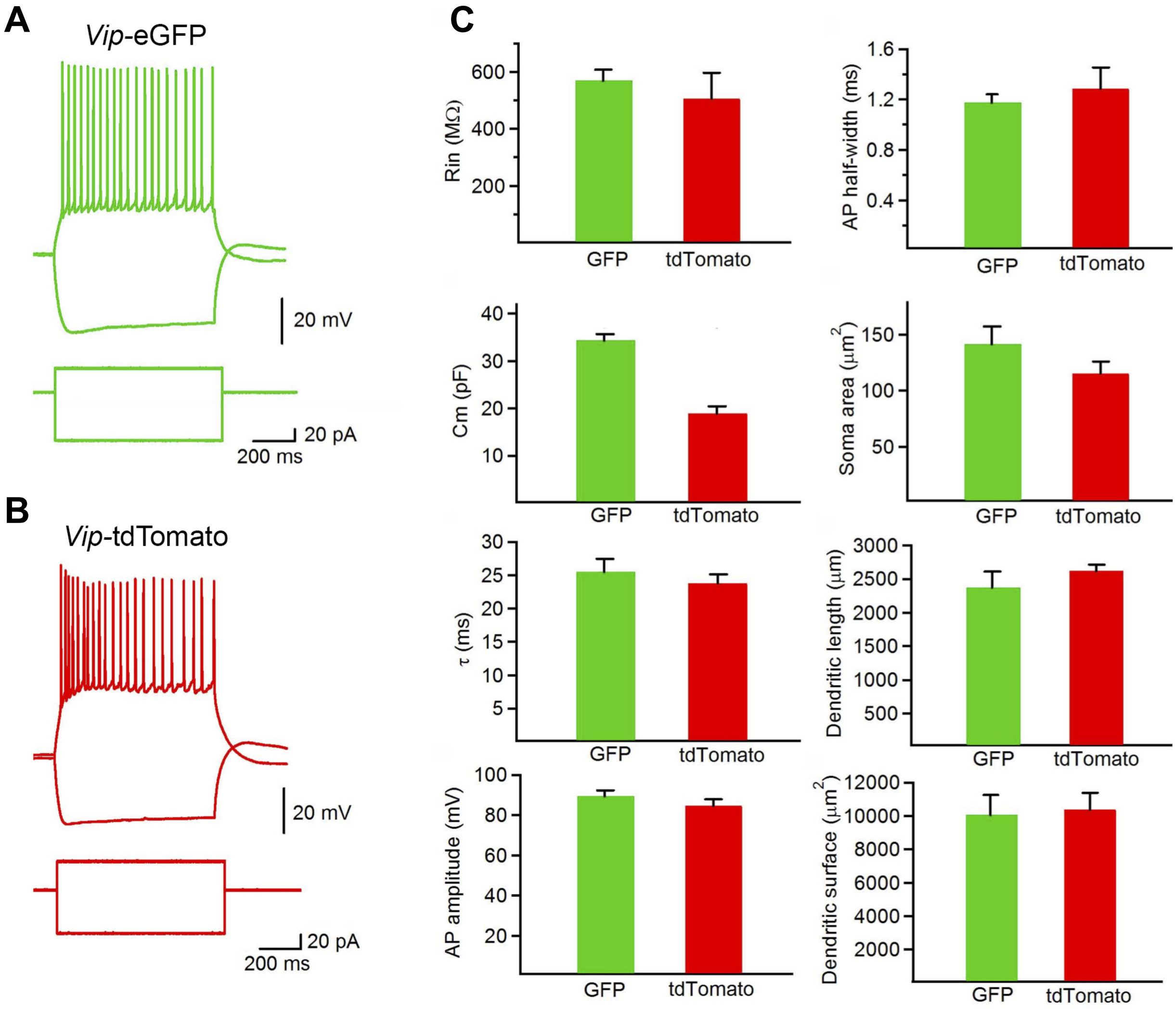
A
Vip-eGFP
20 mV
20 pA
200 ms
B
Vip-tdTomato
20 mV
20 pA
200 ms
C
Rin (MΩ)
600
400
200
GFP
tdTomato
AP half-width (ms)
1.6
1.2
0.8
0.4
GFP
tdTomato
Cm (pF)
40
30
20
10
GFP
tdTomato
Soma area (μm2)
150
100
50
GFP
tdTomato
τ (ms)
30
25
20
15
10
5
GFP
tdTomato
Dendritic length (μm)
3000
2500
2000
1500
1000
500
GFP
tdTomato
AP amplitude (mV)
100
80
60
40
20
GFP
tdTomato
Dendritic surface (μm2)
12000
10000
8000
6000
4000
2000
GFP
tdTomato

**Figure S2.**

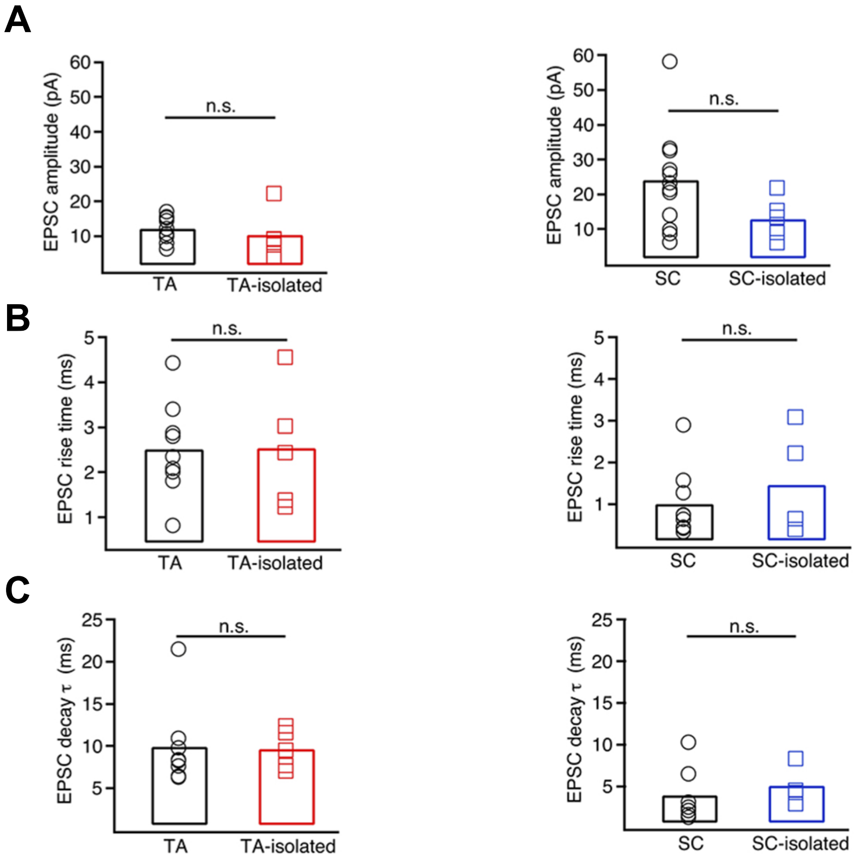
A
EPSC amplitude (pA)
n.s.
TA
TA-isolated
SC
SC-isolated
B
EPSC rise time (ms)
C
EPSC decay τ (ms)

**Figure S3.**

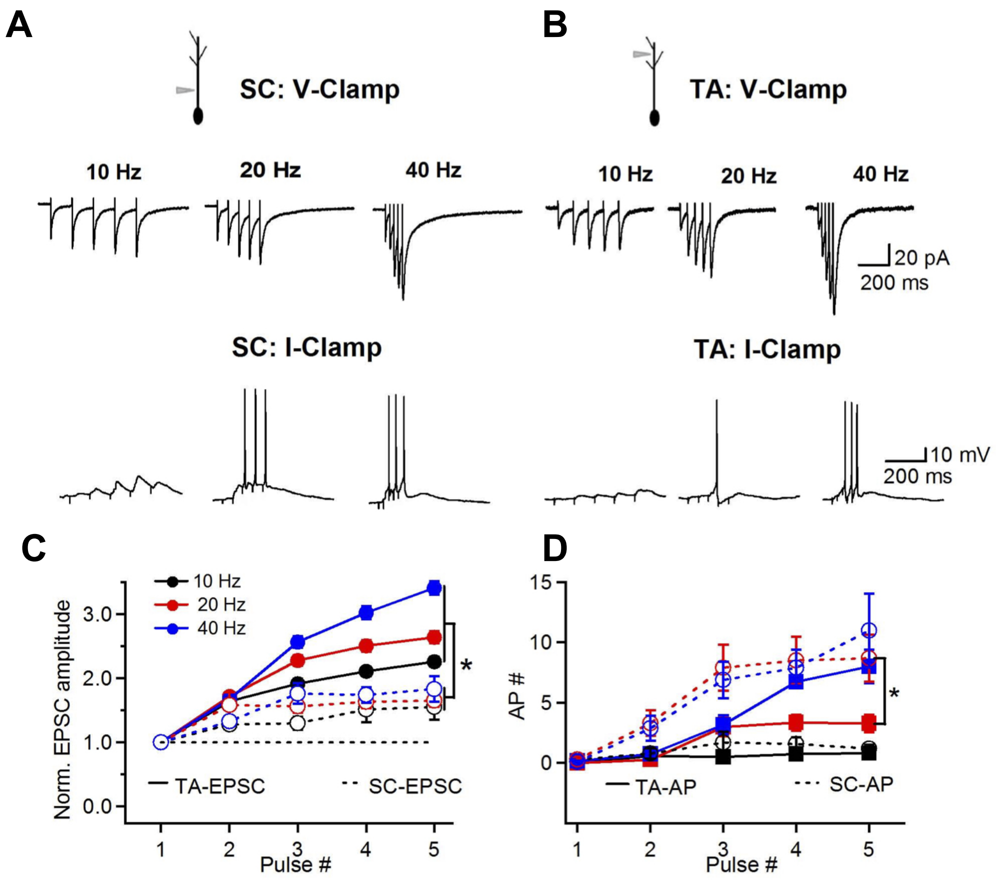
A
SC: V-Clamp
10 Hz
20 Hz
40 Hz
SC: I-Clamp
B
TA: V-Clamp
10 Hz
20 Hz
40 Hz
20 pA
200 ms
TA: I-Clamp
10 mV
200 ms
C
10 Hz
20 Hz
40 Hz
Norm. EPSC amplitude
TA-EPSC
SC-EPSC
Pulse #
D
AP #
TA-AP
SC-AP
Pulse #

**Figure S4.**

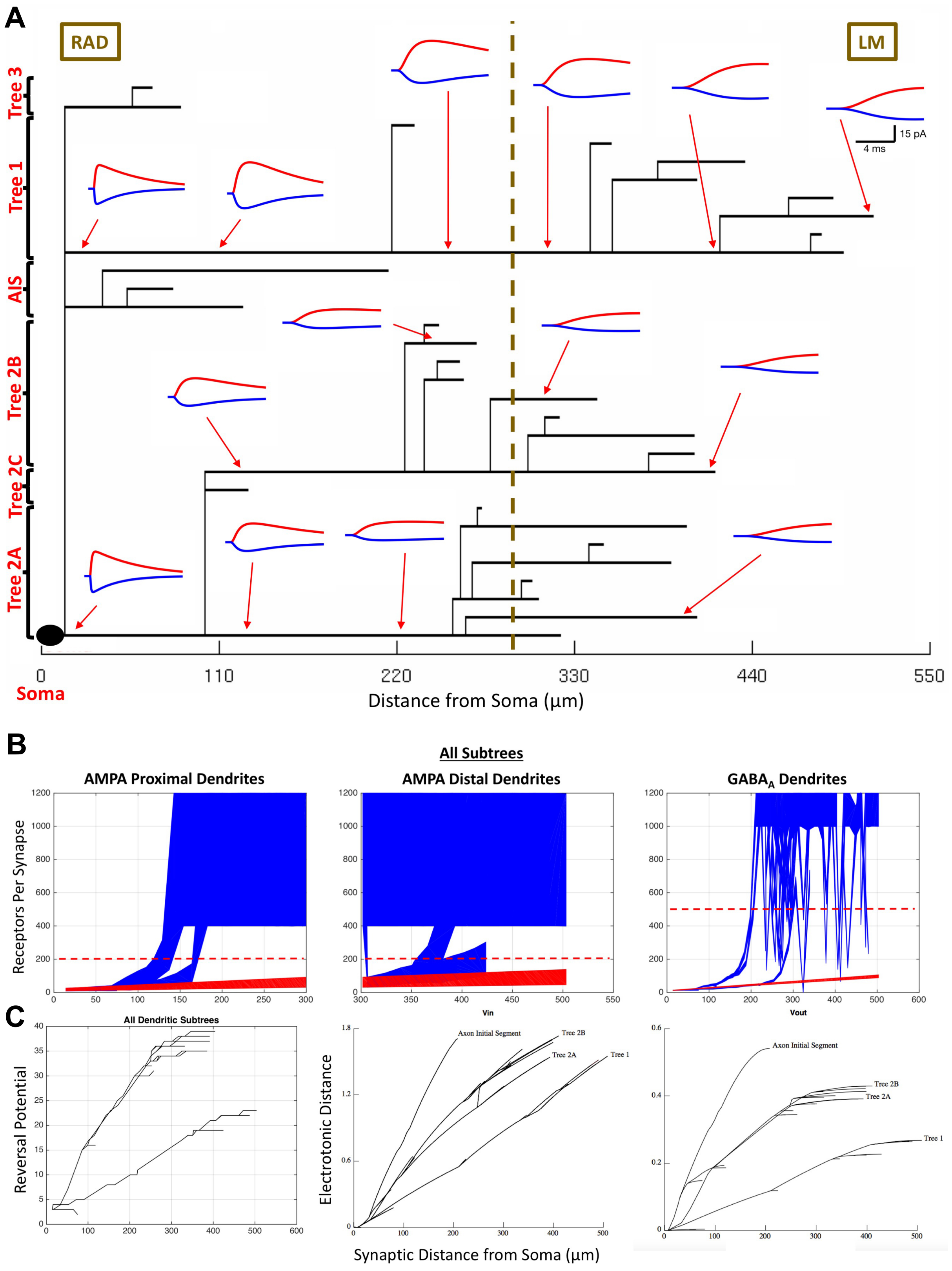
A
RAD
LM
Tree 3
Tree 1
AIS
Tree 2B
Tree 2C
Tree 2A
15 pA
4 ms
0
110
220
330
440
550
Soma
Distance from Soma (μm)
B
All Subtrees
AMPA Proximal Dendrites
AMPA Distal Dendrites
GABAA Dendrites
Receptors Per Synapse
Vin
Vout
C
All Dendritic Subtrees
Reversal Potential
Electrotonic Distance
Axon Initial Segment
Tree 2B
Tree 2A
Tree 1
Synaptic Distance from Soma (μm)

**Figure S5.**

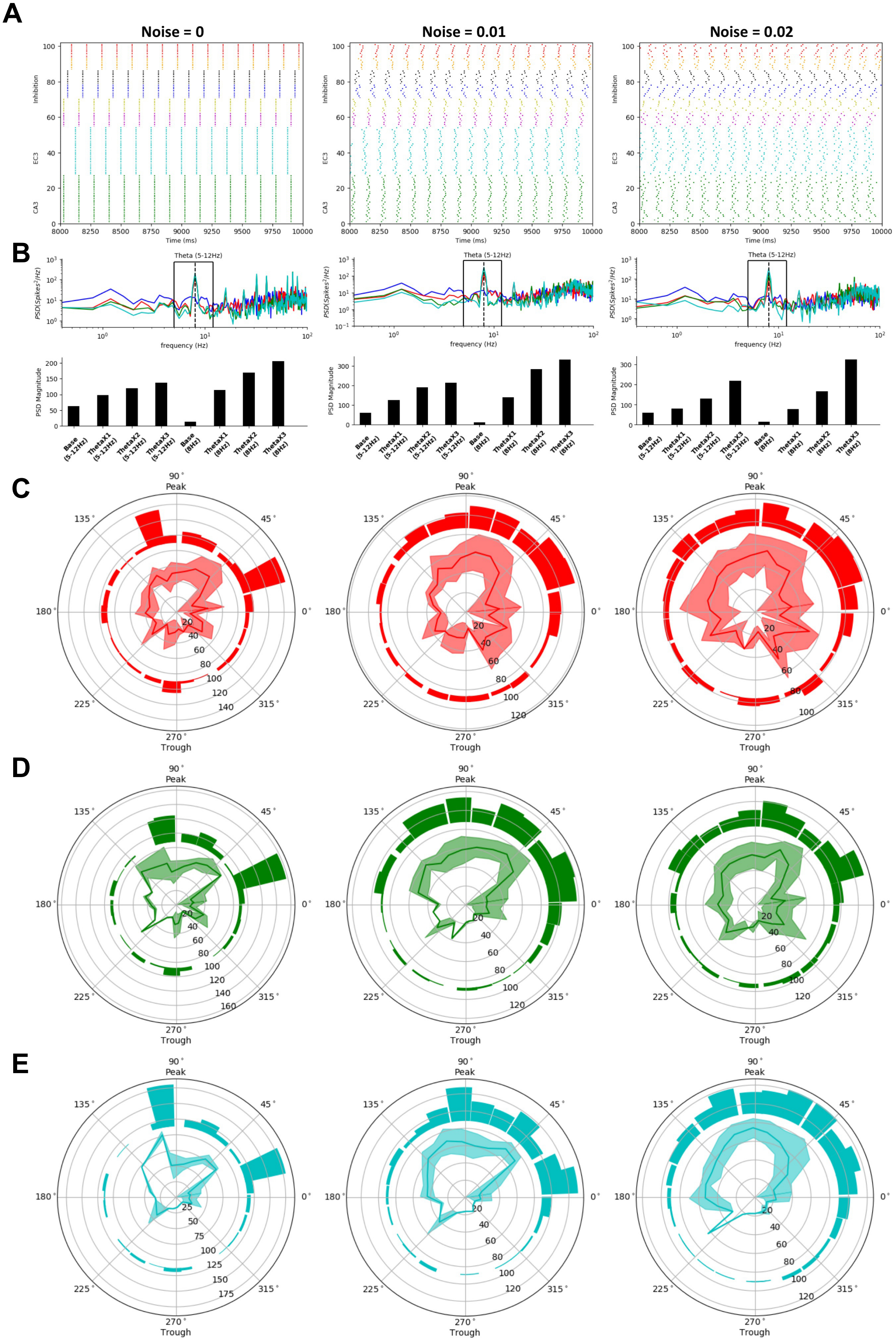
A
Noise = 0
Noise = 0.01
Noise = 0.02
Inhibition
EC3
CA3
Time (ms)
B
Theta (5-12Hz)
frequency (Hz)
PSD Magnitude
Base (5-12Hz)
ThetaX1 (5-12Hz)
ThetaX2 (5-12Hz)
ThetaX3 (5-12Hz)
Base (8Hz)
ThetaX1 (8Hz)
ThetaX2 (8Hz)
ThetaX3 (8Hz)
C
D
E
Peak
Trough

**Figure S6.**

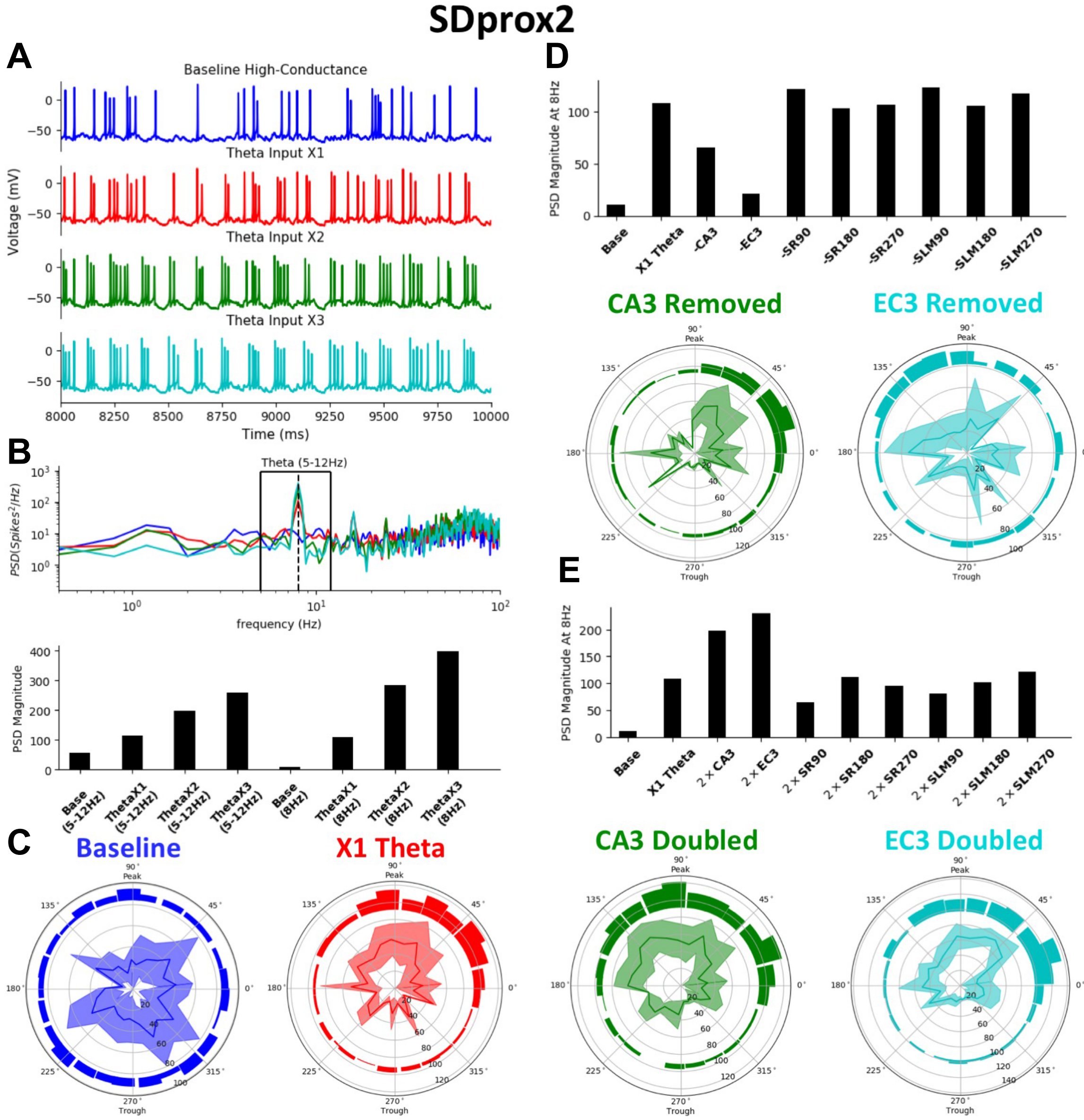
SDprox2
A
Baseline High-Conductance
Theta Input X1
Theta Input X2
Theta Input X3
Voltage (mV)
Time (ms)
B
Theta (5-12Hz)
frequency (Hz)
PSD Magnitude
Base (5-12Hz)
ThetaX1 (5-12Hz)
ThetaX2 (5-12Hz)
ThetaX3 (5-12Hz)
Base (8Hz)
ThetaX1 (8Hz)
ThetaX2 (8Hz)
ThetaX3 (8Hz)
C
Baseline
X1 Theta
D
PSD Magnitude At 8Hz
Base
X1 Theta
-CA3
-EC3
-SR90
-SR180
-SR270
-SLM90
-SLM180
-SLM270
CA3 Removed
EC3 Removed
E
PSD Magnitude At 8Hz
Base
X1 Theta
2×CA3
2×EC3
2×SR90
2×SR180
2×SR270
2×SLM90
2×SLM180
2×SLM270
CA3 Doubled
EC3 Doubled
Peak
Trough

**Figure S7.**

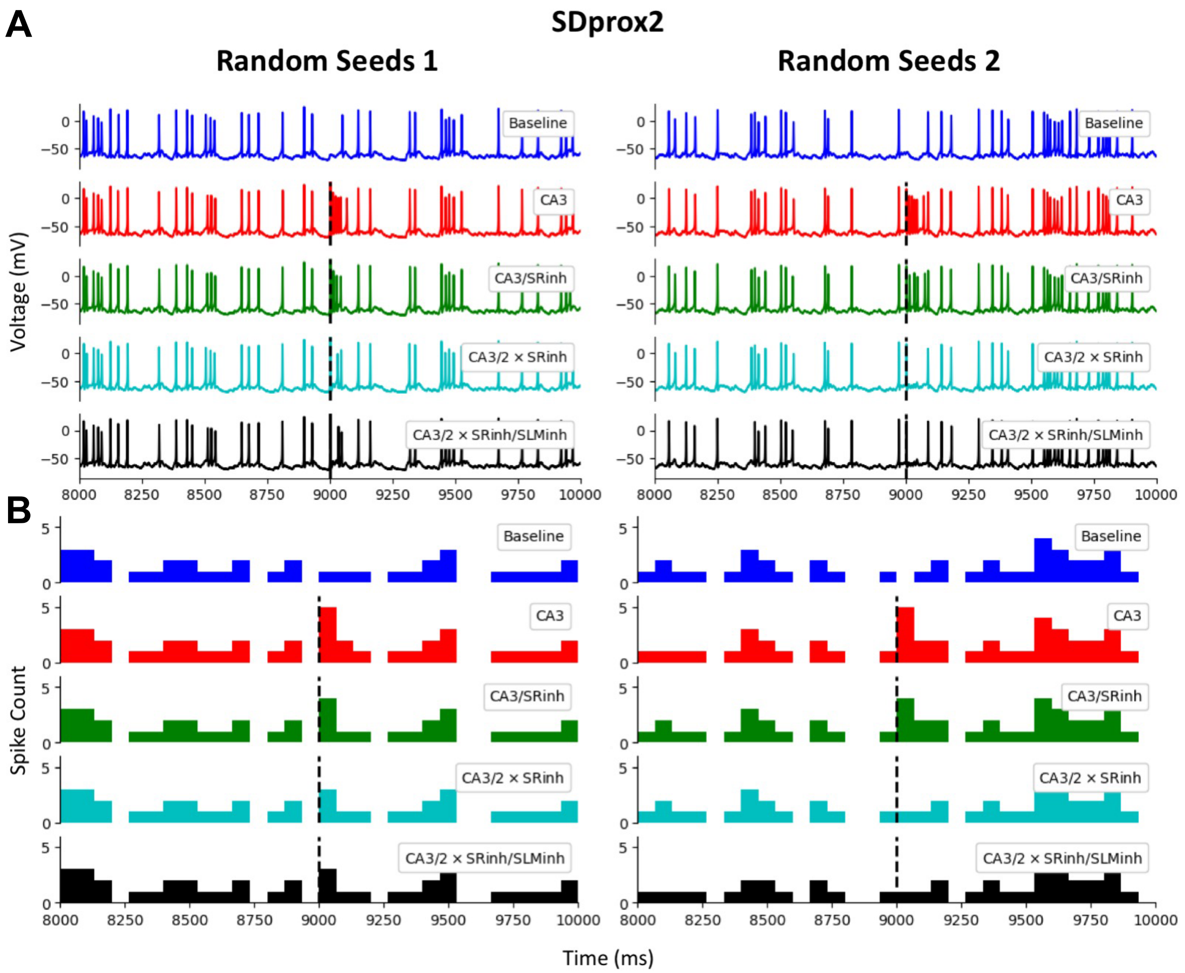
A
SDprox2
Random Seeds 1
Random Seeds 2
Voltage (mV)
Baseline
CA3
CA3/SRinh
CA3/2 × SRinh
CA3/2 × SRinh/SLMinh
B
Spike Count
Time (ms)

**Figure S8.**

**Post hoc *in vitro***

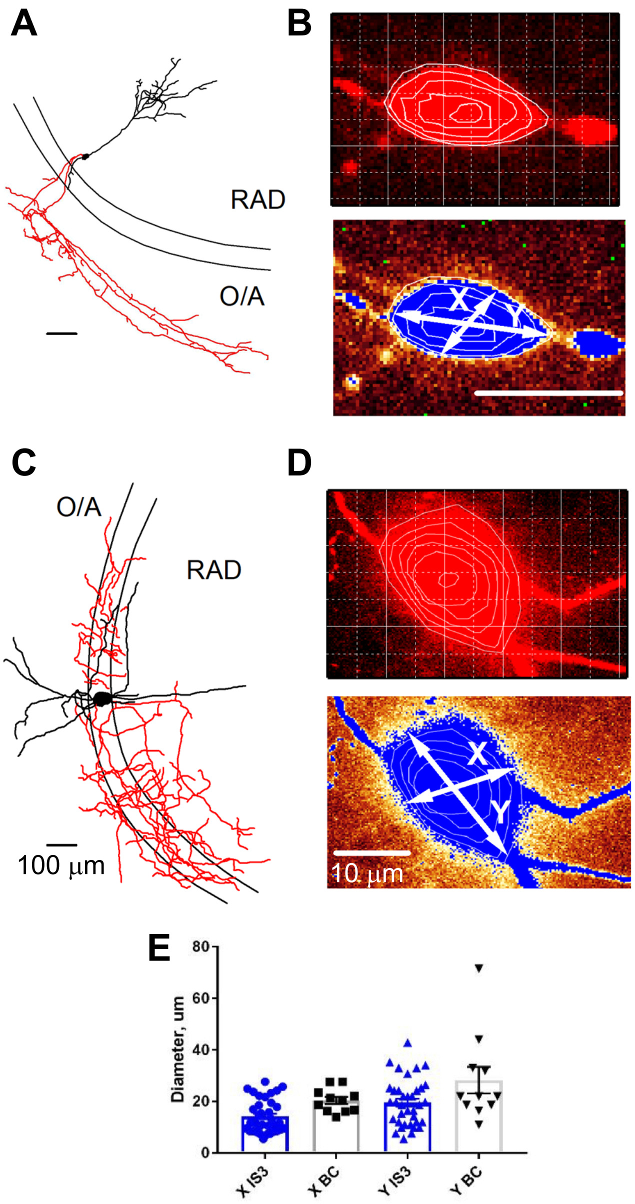


**Figure S9.**

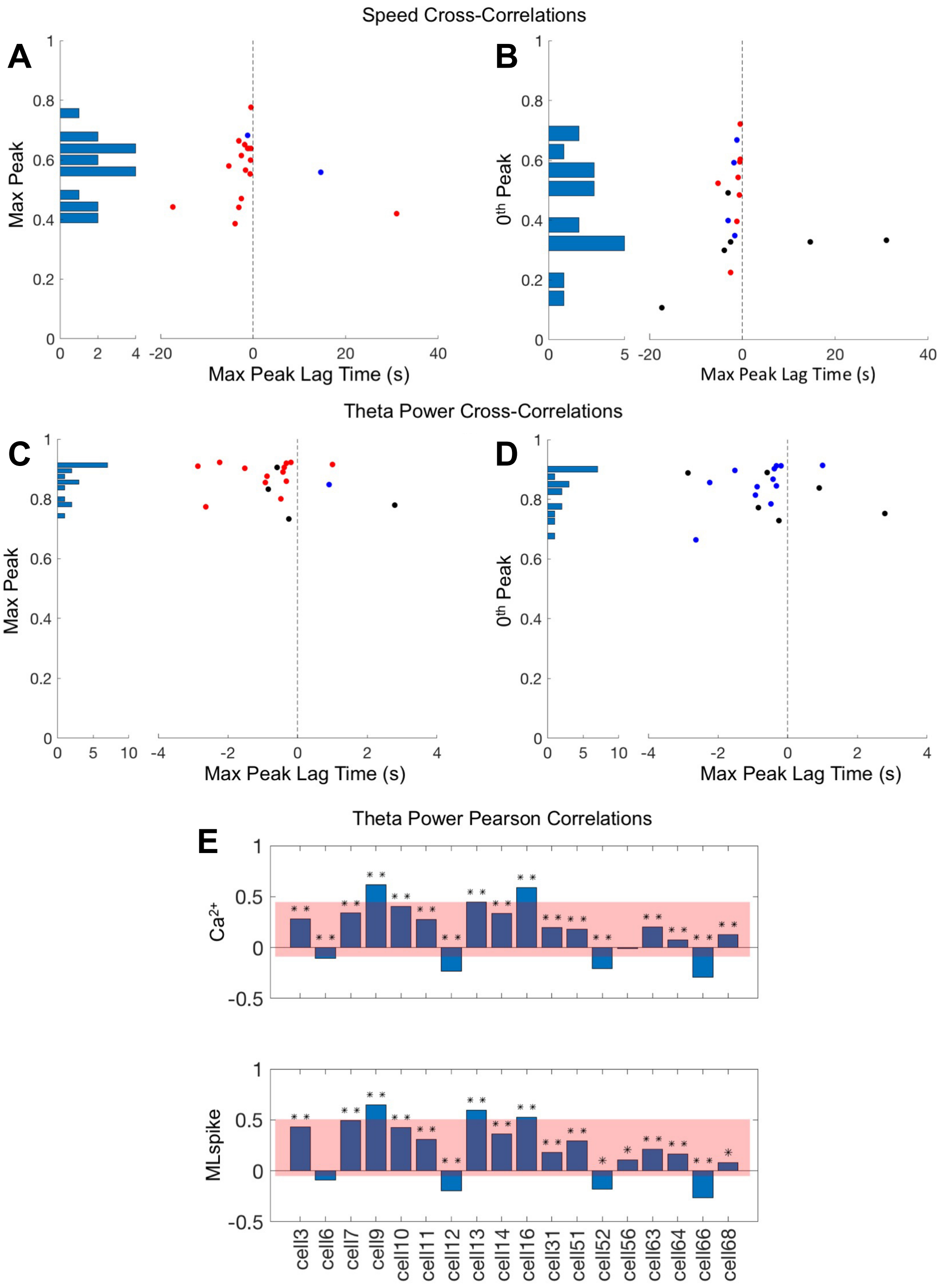

Speed Cross-Correlations
A
B
Max Peak
0th Peak
Max Peak Lag Time (s)
Theta Power Cross-Correlations
C
D
Theta Power Pearson Correlations
E
Ca2+
MLspike
cell3
cell6
cell7
cell9
cell10
cell11
cell12
cell13
cell14
cell16
cell31
cell51
cell52
cell56
cell63
cell64
cell66
cell68


**Figure S10.**

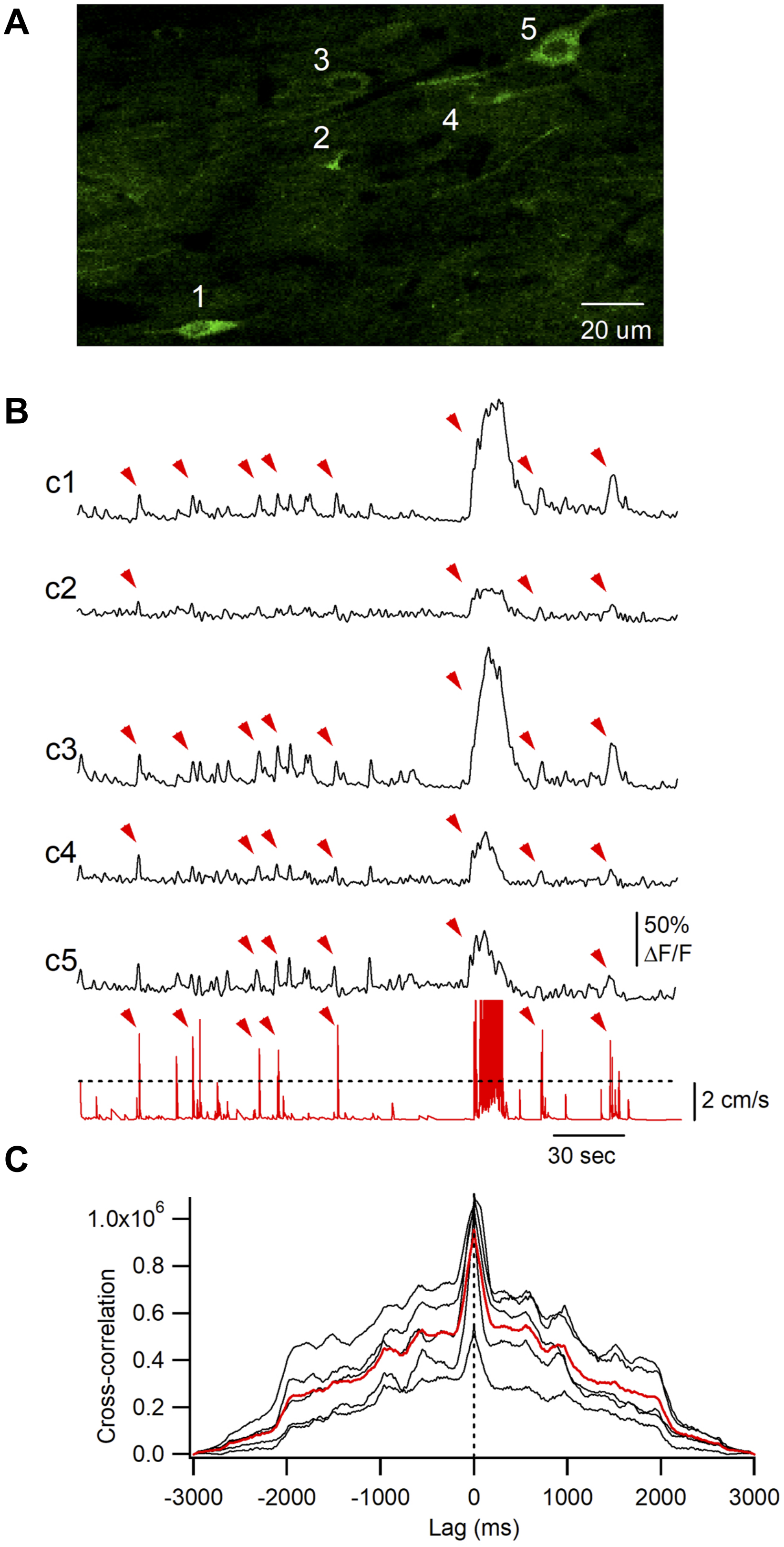
A
5
3
4
2
1
20 um
B
c1
c2
c3
c4
c5
50%
ΔF/F
2 cm/s
30 sec
C
1.0x10^6
0.8
0.6
0.4
0.2
0.0
Cross-correlation
-3000
-2000
-1000
0
1000
2000
3000
Lag (ms)

**Figure S11.**

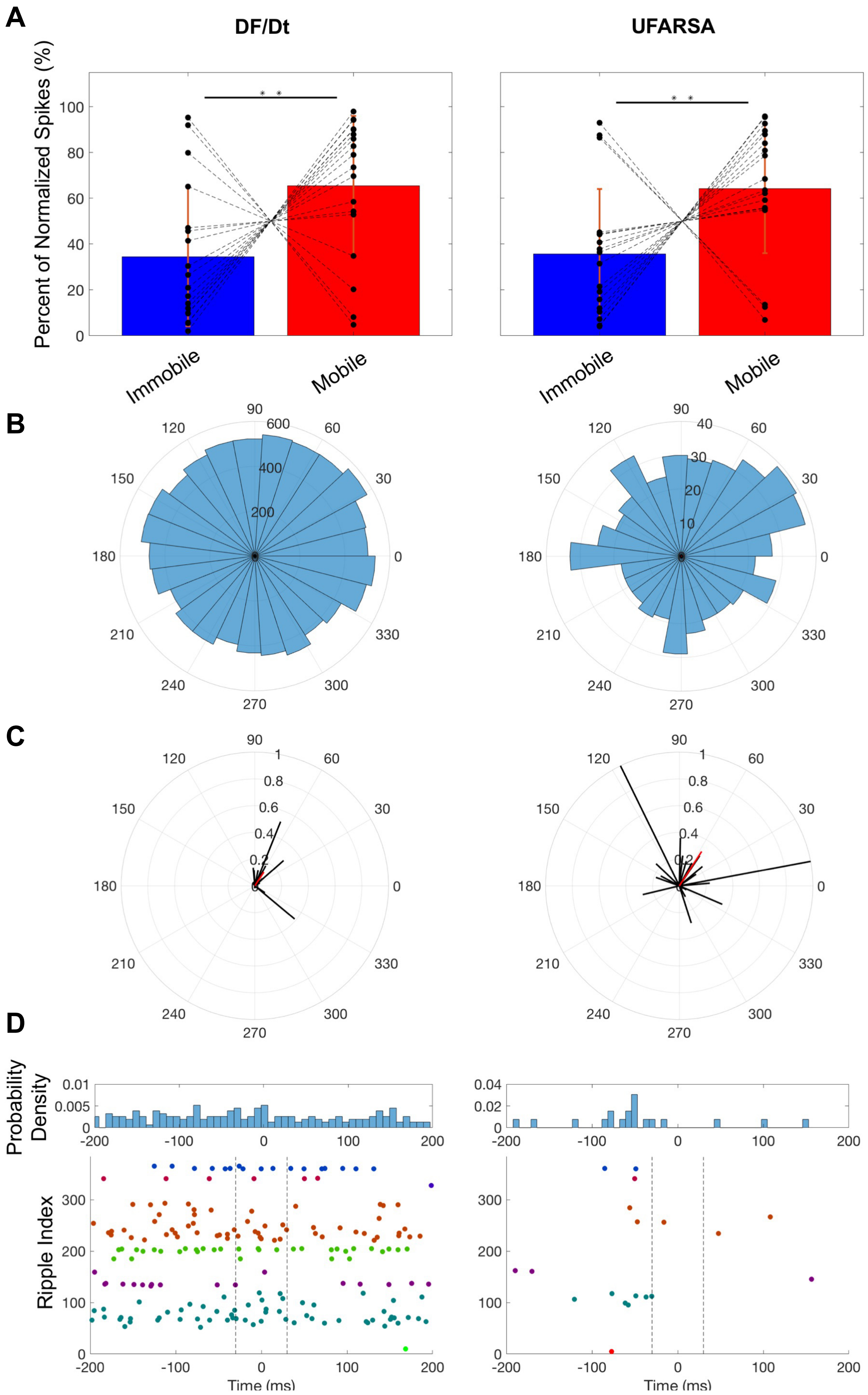
A
DF/Dt
UFARSA
Percent of Normalized Spikes (%)
Immobile
Mobile
B
C
D
Probability Density
Ripple Index
Time (ms)

**Figure S12.**
